# Pan-cancer analysis reveals mtDNA copy number as a key determinant of mutational load and disease progression in cancer

**DOI:** 10.1101/2025.11.18.689019

**Authors:** Anamika Acharyya, Riddhiman Dhar

**Affiliations:** Department of Bioscience and Biotechnology, IIT Kharagpur, India

**Keywords:** MtDNA copy number, Pan-cancer, Mutational load, Differential expression, Cancer progression, Chemotherapy response

## Abstract

Mitochondrial DNA (mtDNA) copy number determines the functional state of mitochondria and thus, influences cellular energy production, growth, metabolism, and stress response. Variation in mtDNA copy number has been observed across many cancer types and has been linked to changes in gene expression programs. However, whether mtDNA copy number has any influence on mutation accumulation in cancer is unknown. Further, the actual impact of mtDNA copy number variation on cancer progression remains unclear with conflicting reports across a few cancer types. Here, through a pan-cancer analysis of whole genome data, we show that mtDNA copy number increases with an increase in mutational load in cancer. The increase in mtDNA copy number bears a signature of compensation for detrimental effects on mitochondrial function caused by increased mutational load. We also show that the samples with low mtDNA generally have increased expression of cancer promoting genes whereas high mitochondrial activity is linked to higher activity of tumor suppressor genes. We further demonstrate that low mtDNA copy number increases the likelihood of chemotherapy resistance. Taken together, these results reveal a central role of mtDNA copy number variation in determining mutational load, disease progression, and therapy response across a variety of cancer types. These findings can help design new strategies for disease management and therapy development in cancer.

## Introduction

Mitochondria have a central role in vital cellular processes including energy generation, metabolism, respiration, iron homeostasis, stress response, and apoptosis (McBride et al., 2006). Proper functioning of mitochondria requires ∼1500 proteins in human cells (Wallace, 2005), majority of which are encoded by the nuclear DNA. In addition, mitochondria harbour their own DNA molecules (mtDNA) in multiple copies encoding for 13 proteins in human cells that are essential for mitochondrial function (Wallace, 2005). Thus, mutations in the nuclear DNA or the mtDNA and changes in the mtDNA copy number can alter mitochondrial function and thereby, can affect many important cellular processes.

The role of mitochondria in cancer has received widespread attention over the years (Wallace, 2012; Altier, 2023). One of the hallmarks of cancer includes switching to aerobic glycolysis (the Warburg effect) from cellular respiration for energy generation (Warburg, 1956). In addition, tumor cells show several other metabolic changes - such as increased requirement of glutamine and synthesis of nucleotide precursor molecules - that are associated with mitochondrial function (Ahn and Metallo, 2015; Li et al., 2024). These metabolic changes can result from mutations in genes associated with mitochondrial function and changes in mtDNA copy number and thus, there has been a major push towards identifying mtDNA mutations and mapping mtDNA copy number alterations across cancer types (Reznik et al., 2016; Yuan et al., 2020).

Earliest studies have highlighted changes in mtDNA copy number in specific and limited number of cancer types and observed tumor-specific changes in copy number (Mambo et al., 2005; Wang et al., 2005; Marucci et al., 2013; Wu et al., 2005; Cui et al., 2013). Development of The Cancer Genome Atlas (TCGA) and the International Cancer Genome Consortium (ICGC) datasets, with availability of large-scale whole genome and whole exome sequencing data, enabled systematic investigation of the changes in mtDNA copy number across a variety of cancer types (Reznik et al., 2016; Yuan et al., 2020). These studies revealed tumor-specific nature of changes in mtDNA copy number but differed in their quantification of these changes, likely due to use of different type of data. For example, Reznik et al. (2016), using whole exome and whole genome data from TCGA for mapping mtDNA copy number changes, observed a significant decrease in mtDNA copy number in kidney, breast, liver, and head & neck cancer and a significant increase in lung adenocarcinoma. In comparison, Yuan et al. (2020), using only whole genome data of the Pan-Cancer Analysis of Whole Genomes (PCAWG) dataset, also observed a significant decrease in mtDNA copy number in Kidney renal cell carcinoma and liver hepatocellular carcinoma, but observed a significant increase in copy number in pancreatic adenocarcinoma, lung squamous cell carcinoma, and chronic lymphocytic leukaemia.

Despite the large-scale quantification of mtDNA copy number variation in cancer, a key question on whether mtDNA copy number variation has any association with the mutation accumulation process in cancer genomes remains unanswered. This is of particular interest given that an earlier study revealed an association between occurrence of mtDNA Copy number and mutations in the *IDH1*, *IDH2* and *PTEN* genes in low grade glioma (Reznik et al., 2016). In addition, it is not clear whether changes in mtDNA copy number, through changes in gene expression and cellular processes, can influence the rate of occurrence and signatures of mutations in cancer, or whether mtDNA copy number alteration is merely a byproduct of mutations occurring in cancer cells. Intriguingly, in kidney chromophobe carcinoma, cancer samples with more mtDNA mutations had increased mtDNA copy number (Reznik et al., 2016), hinting at a possible link between increased mtDNA copy number and mtDNA mutational load in these samples.

In addition, how variation in mtDNA copy number influences cancer progression remains unclear, with several conflicting reports from studies on a limited number of cancer types (Wang et al., 2006; Lin et al., 2010). Two different studies on colorectal cancer reached contradictory conclusions regarding the association of mtDNA copy number with cancer progression (Cui et al., 2013; Sun et al., 2018). In addition, low mtDNA copy number was linked to cancer progression and decreased survival of patients in ovarian cancer (Wang et al., 2006), and hepatocellular carcinoma (Yamada et al., 2006), whereas high mtDNA copy number was linked to cancer progression and poor prognosis in head & neck cancer (Kim et al., 2004) and esophageal squamous cell cancer (Lin et al., 2010).

Further, the molecular processes by which mtDNA copy number variation can influence cancer progression remains poorly understood. As expected, correlation of gene expression with mtDNA copy number (Reznik et al., 2016; Reznik et al., 2017) and weighted gene co-expression network analysis (Yuan et al., 2020) revealed changes in genes associated with oxidative phosphorylation, mitochondrial metabolism, DNA repair, and cell cycle. However, these results did not investigate whether the changes in these processes could potentially facilitate or inhibit cancer progression. Intriguingly, in patients with shorter survival across cancer types, oxidative phosphorylation was downregulated and was correlated with induction of epithelial-mesenchymal pathway (Gaude and Frezza, 2016), suggesting an association between mitochondrial function and cancer progression.

In this work, through a pan-cancer analysis of mtDNA copy number and mutations from PCAWG dataset, we observed a positive correlation between mtDNA copy number in cancer samples and their mutational load. We also observed a signature of compensation for mitochondrial functions associated with mtDNA copy number increase. Specifically, across different cancer types, mtDNA copy number showed positive correlation with the number of mitochondria-localized genes containing truncating, nonsynonymous or intron/splice site/UTR mutations. In addition, differential expression analysis revealed upregulation of cancer promoting genes in low mtDNA samples and upregulation of suppressor genes in high mtDNA samples in several cancer types. Further, mitochondrial gene expression could predict activity of the p53 pathway and patients with low mtDNA copy number had an increased likelihood of developing chemoresistance. Taken together, these results highlight a central role of mitochondria in cancer progression which could be exploited for devising new cancer management and treatment strategies.

## Results

### mtDNA copy number is positively correlated with mutational load in cancer samples

To test whether mtDNA copy number has any association with occurrence of mutations in cancer (Fig. 1A), we obtained the single nucleotide variant and indel data from PCAWG dataset along with the mtDNA copy number data from The Cancer Mitochondria Atlas (TCMA) dataset. In addition, we also included the data on single nucleotide variants and indels in mtDNA that were mapped in the TCMA dataset.

**Figure 1.**
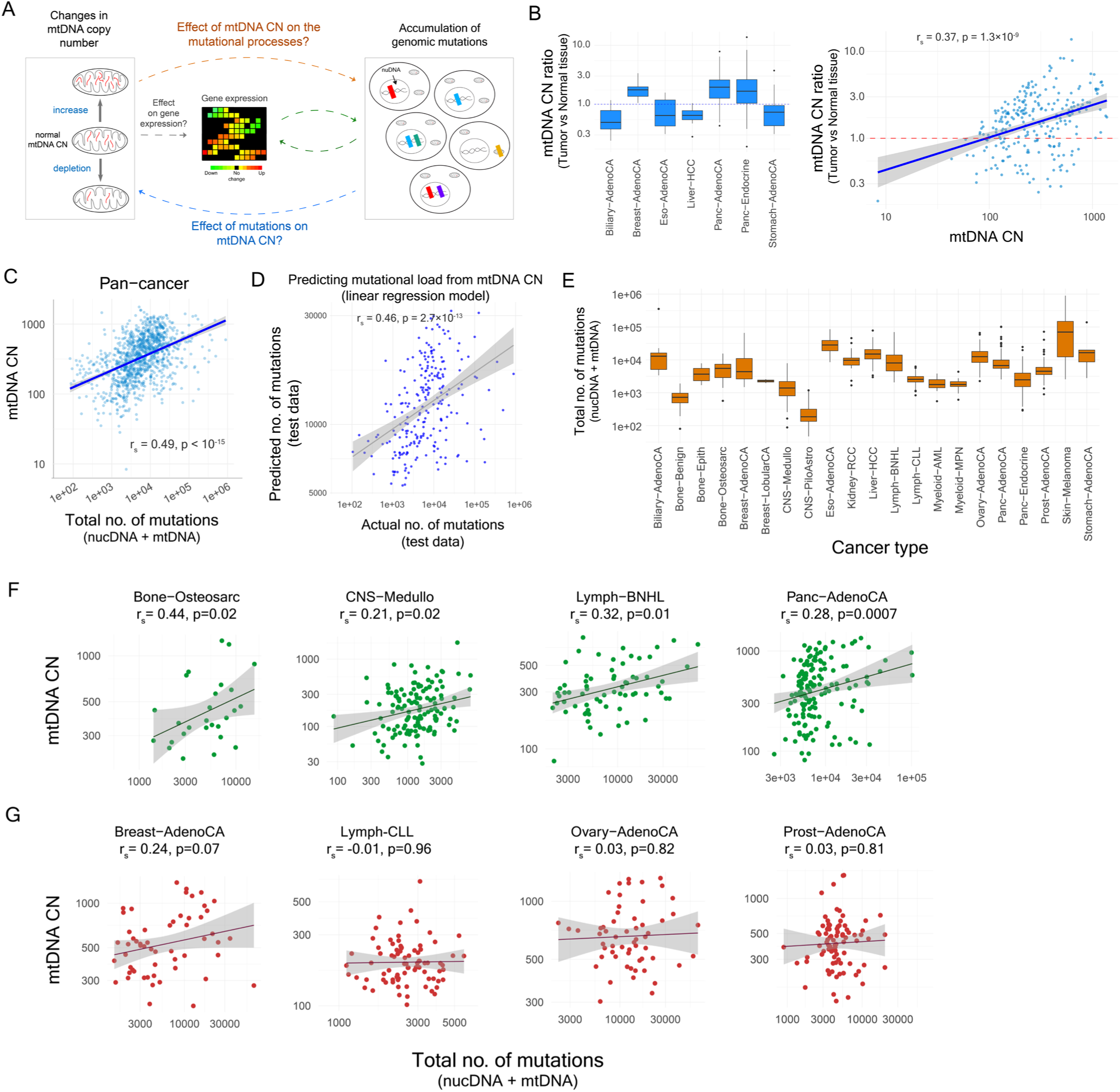
MtDNA copy number increases with an increase in mutational load in cancer samples. **(A)** Schematic diagram depicting the key question - whether mtDNA copy number has any association with the mutational processes in cancer. **(B)** The ratio of mtDNA copy number between cancer and matched normal tissue samples. **(C)** Correlation between mtDNA copy number and the ratio of mtDNA copy number between cancer and matched normal tissue samples. **(D)** Pan-cancer correlation between the total number of mutations (nuclear DNA + mtDNA mutations) and mtDNA copy number. **(E)** The distributions of the total number of mutations in patient samples across 21 cancer types. **(F-G)** Correlation between the total number of mutations and mtDNA copy number in individual cancer types. (F) shows the cancer types where significant correlation (p<0.05) exist and (G) shows the cancer types where there is no significant correlation. The solid lines in scatter plots show linear fits to the data and the shaded grey regions show 95% confidence intervals. Spearman’s correlation coefficient and corresponding p-values are shown.

We calculated the ratio of mtDNA copy number in cancer and matched normal tissue samples for seven cancer types for which at least three matched normal samples were available (Fig. 1B). The mtDNA copy number in cancer sample was significantly correlated with the ratio of mtDNA copy number between cancer and normal samples in these cancer types (Spearman’s correlation coefficient r_s_=0.37, p=1.3×10^-9^; Fig. 1B). Similarly, we also saw significant correlation between mtDNA copy number in cancer samples and the ratio of mtDNA copy number between cancer and matched normal blood samples (r_s_=0.74, p<10^-15^; Fig. S1). Thus, we used the absolute tumor mtDNA copy number as a marker for the change of mtDNA copy number in tumor compared to normal samples for all subsequent analyses. We also used samples with tumor purity of at least 0.5, to ensure that the inferences obtained were not largely due to presence of adjacent normal tissue. In total, we had a total of 1230 samples across 24 cancer types (Table S1).

We first tested the correlation between mtDNA copy number and the total number of mutations in cancer genomes – comprising of both nuclear DNA and mtDNA single nucleotide variants (SNVs) and small indels. In pan-cancer analysis, the mtDNA copy number showed significant positive correlation with the total number of mutations (r_s_=0.49, p<10^-15^; Fig. 1C). To test whether mtDNA copy number alone could predict mutational load of cancer samples, we also built a predictive linear regression model where we fit the linear regression on the randomly sampled training data (80% of the full data) and predicted the total number of mutations from mtDNA copy number in test data (remaining 20% of the full data). The predicted number of mutations was significantly correlated with the actual number of mutations in the test samples (r_s_=0.46, p = 2.7×10^-13^, Fig. 1D), suggesting that the mtDNA copy number could well predict the extent of mutational load in cancer samples.

In pan-cancer analysis, we also observed significant correlation of mtDNA copy number with the number of nuclear DNA SNVs (r_s_=0.47, p<10^-15^), nuclear DNA indels (r_s_=0.50, p<10^-15^), and mtDNA SNVs (r_s_=0.25, p=1.5×10^-13^) (Fig. S2A-C). However, we did not see any significant correlation of mtDNA copy number with the number of mtDNA indels (r_s_=0.03, p=0.57; Fig. S2D). In addition, we did not see any correlation between the number of mutations – comprising of both SNVs and indels - in nuclear DNA and the number of mutations in mtDNA across samples (r_s_=0.04, p=0.56; Fig. S3).

As we observed substantial variation in mtDNA copy number and total number of mutations in different cancer types (Fig. 1E and Fig. S4), we also tested for correlation between these two parameters in individual cancer types. We observed significant correlation in pancreatic adenocarcinoma (r_s_=0.28, p=0.0007), Lymph-BNHL (r_s_=0.32, p=0.01), bone-osteosarcoma (r_s_=0.44, p=0.02), and CNS medulloblastoma (r_s_=0.21, p=0.02) (Fig. 1F). But we did not find significant correlation in other cancer types such as breast adenocarcinoma, prostate adenocarcinoma, and kidney-RCC (Fig. 1G and Fig. S5).

### Tumor ploidy and nuclear indels are top predictors of mtDNA copy number

To further investigate the molecular causes that could drive the changes in mtDNA copy number in cancer samples, we built predictive models using linear regression and random forest framework. We considered a total of 87 molecular features derived for these samples (Table S2). The features ranged from total number of mutations, number of different type of mutations (SNVs, indels), number of nuclear DNA and mtDNA mutations, percentage of specific types of nucleotide changes and indels, to the number of genes affected by truncating (nonsense and frameshift indels), non-truncating (missense, in-frame indels) and silent mutations, as well as the number of affected genes associated with mitochondrial function (Table S2).

We used linear regression and random forest framework which can capture linear and non-linear associations, respectively, between the molecular features and mtDNA copy number. Since several molecular features in our data are likely to be highly correlated with each other (multi-collinearity), we used ridge and lasso regression models for linear regression (see Methods; Fig. 2A). We randomly split the full data into training (80% data) and test (remaining 20% data) sets, fit the models to the training data, and then predicted mtDNA copy number on the test data, and in the process, also identifying the key molecular features (Fig. 2A). Predicted mtDNA copy number from ridge, lasso and random forest models showed significant correlation with the actual mtDNA copy number, out of which the random forest model showed slightly better predictive capability (Ridge: r_s_ = 0.59, p<10^-15^; Lasso: r_s_ = 0.58; p<10^-15^; Random Forest: r_s_ = 0.67, p<10^-15^; Fig. 2B and Fig. S6).

**Figure 2.**
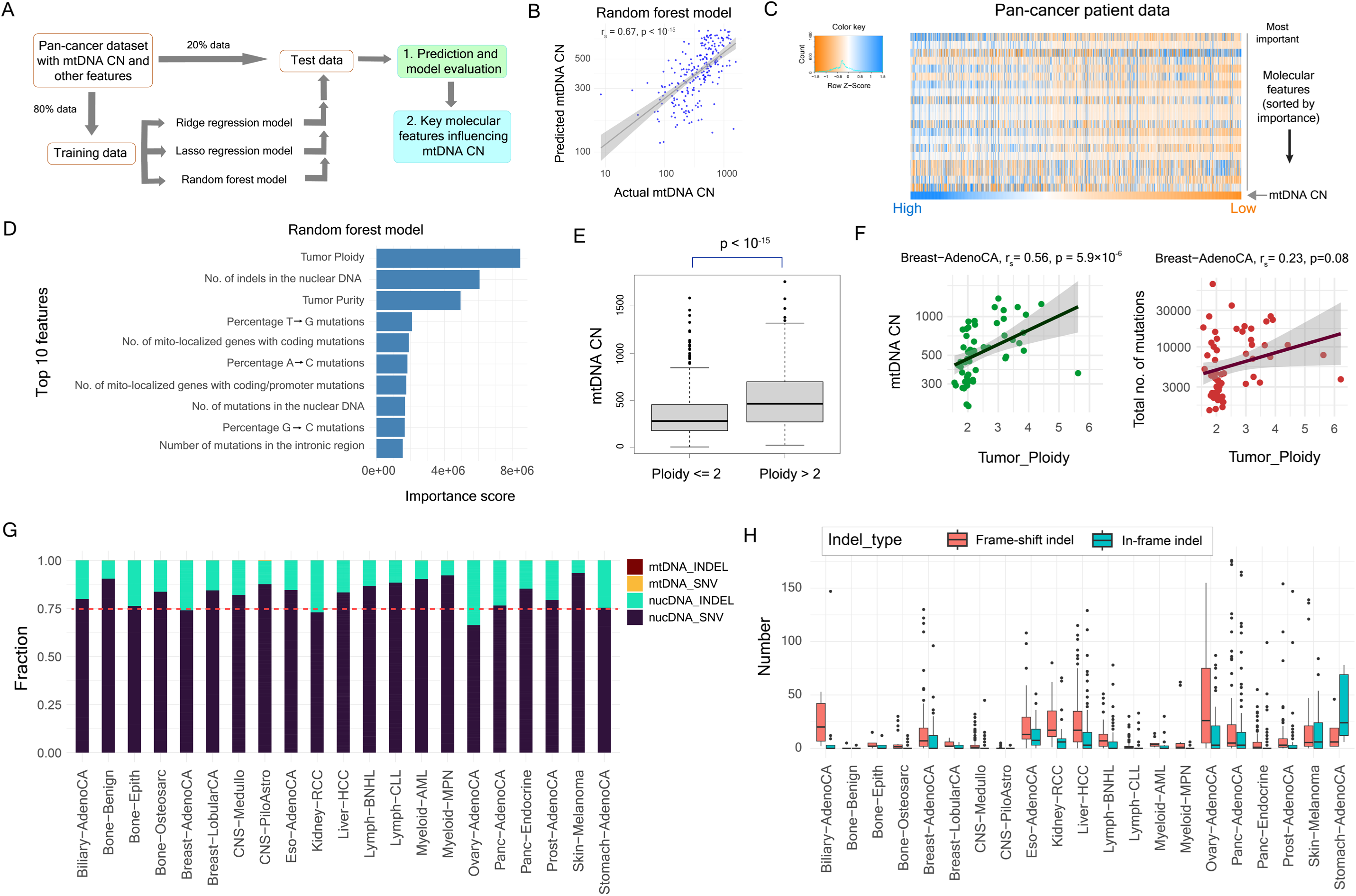
Tumor ploidy and the number of nuclear indels are the top predictors of mtDNA copy number in cancer samples. **(A)** Schematic diagram depicting the framework for predicting mtDNA copy number and for identifying the key molecular features. **(B)** Correlation between the actual mtDNA copy number and the mtDNA copy number predicted by the random forest model. **(C)** Heatmap showing the values of the top 20 features of the random forest model (in rows) in cancer samples (in columns). The samples were sorted according to mtDNA copy number that is also shown in the heatmap. **(D)** The importance scores of the top 10 features from the random forest model. **(E)** Boxplot showing distribution of mtDNA copy number in cancer samples with ploidy ≤ 2 and with ploidy > 2. **(F)** Correlation of tumor ploidy with mtDNA copy number (left) and mutational load (right) in breast adenocarcinoma samples. The solid lines show linear fits to the data. **(G)** Average fraction of nuclear SNVs, indels, and mtDNA SNVs, indels in individual cancer types. **(H)** Distribution of the number of frame-shift and in-frame indels in individual cancer types. In scatter plots, the solid blue lines show linear fits to the data and the shaded grey regions show 95% confidence intervals.

Among the features that were important for prediction of mtDNA copy number (Fig. 2C), tumor ploidy was the top ranked feature across all models (Fig. 2D and Fig. S7A,B). In pan-cancer analysis, samples with tumor ploidy greater than two had significantly higher copies of mtDNA compared to samples with ploidy two or less (Wilcoxon rank sum test, p<10^-15^; Fig. 2E). Similarly, the total number of mutations in samples with tumor ploidy greater than two was significantly higher than the samples with ploidy two or less (p = 3.6×10^-14^; Fig. S8).

Within the individual cancer types, we observed an increase in ploidy among a substantial fraction of samples in adenocarcinoma (breast, esophagus, pancreas, ovarian), bone-osteosarcoma, CNS-medulloblastoma, liver-HCC, pancreatic endocrine and skin melanoma (Fig. S9). However, for some cancer types, even though mtDNA copy number increased significantly with tumor ploidy, the number of mutations did not increase much (liver-HCC) (Fig. 2F and Fig. S10). In lymph-CLL, mtDNA copy number decreased with ploidy and the number of mutations did not show significant increase (Fig. S11A). In prostate adenocarcinoma, the number of mutations increased with ploidy whereas the mtDNA copy number did not change (Fig. S11B). None of these cancer types originally showed positive correlation between mtDNA copy number and mutational load (Fig. 1G and Fig S5).

The number of nuclear indels was the second most important feature in random forest model (Fig. 2D). Similarly, percentage of G deletions was the second most important feature in both Ridge and Lasso regression models (Fig. S7). The average percentage of mtDNA indels was less than 1% in all cancer types, whereas the average percentage of nuclear indels varied between ∼5 to 30% across cancer types. Among all cancer types, samples from Breast-AdenoCA, Kidney-RCC and Ovary-AdenoCA had 25% or more nuclear indels on average (Fig. 2G). In addition, samples from several cancer types had substantially higher numbers of frame-shift indels, that would prematurely truncate proteins and would have significant deleterious effect on gene functions, compared to other cancer types (Fig. 2H). These cancer types included Biliary-AdenoCA, Eso-AdenoCA, Kidney-RCC, Liver-HCC, and Ovary-AdenoCA (Fig. 2H), none of which originally showed correlation between mtDNA copy number and mutational load (Fig. 1D and Fig. S5).

### The increase in mtDNA copy number has a signature of functional compensation

We hypothesized that the positive correlation between mtDNA copy number and mutational load in several cancers was perhaps driven by compensation for loss or reduction in gene function due to increased mutational load. The compensation was also likely to be for loss in mitochondrial functions due to mutations in nuclear and mtDNA encoded genes whose protein products are necessary for mitochondrial functions. Across different cancer types, we observed ∼5-10% such nuclear DNA encoded genes that were affected by mutations (Fig. 3A), and we observed a significant pan-cancer correlation between mtDNA copy number and the number of mitochondria-localized genes with mutations in coding or promoter regions or both (Fig. 3B). Individually, across cancer types, we observed significant correlations between mtDNA copy number and the number of mitochondria-localized genes that contained mutations in coding/promoter regions in two cancer types – namely, breast-adenocarcinoma, and liver-HCC (Fig. 3C) – which originally did not show correlation between mtDNA copy number and mutational load (Fig. 1D and Fig. S5), as well as in four cancer types (Fig. 3D) that originally showed correlation.

**Figure 3.**
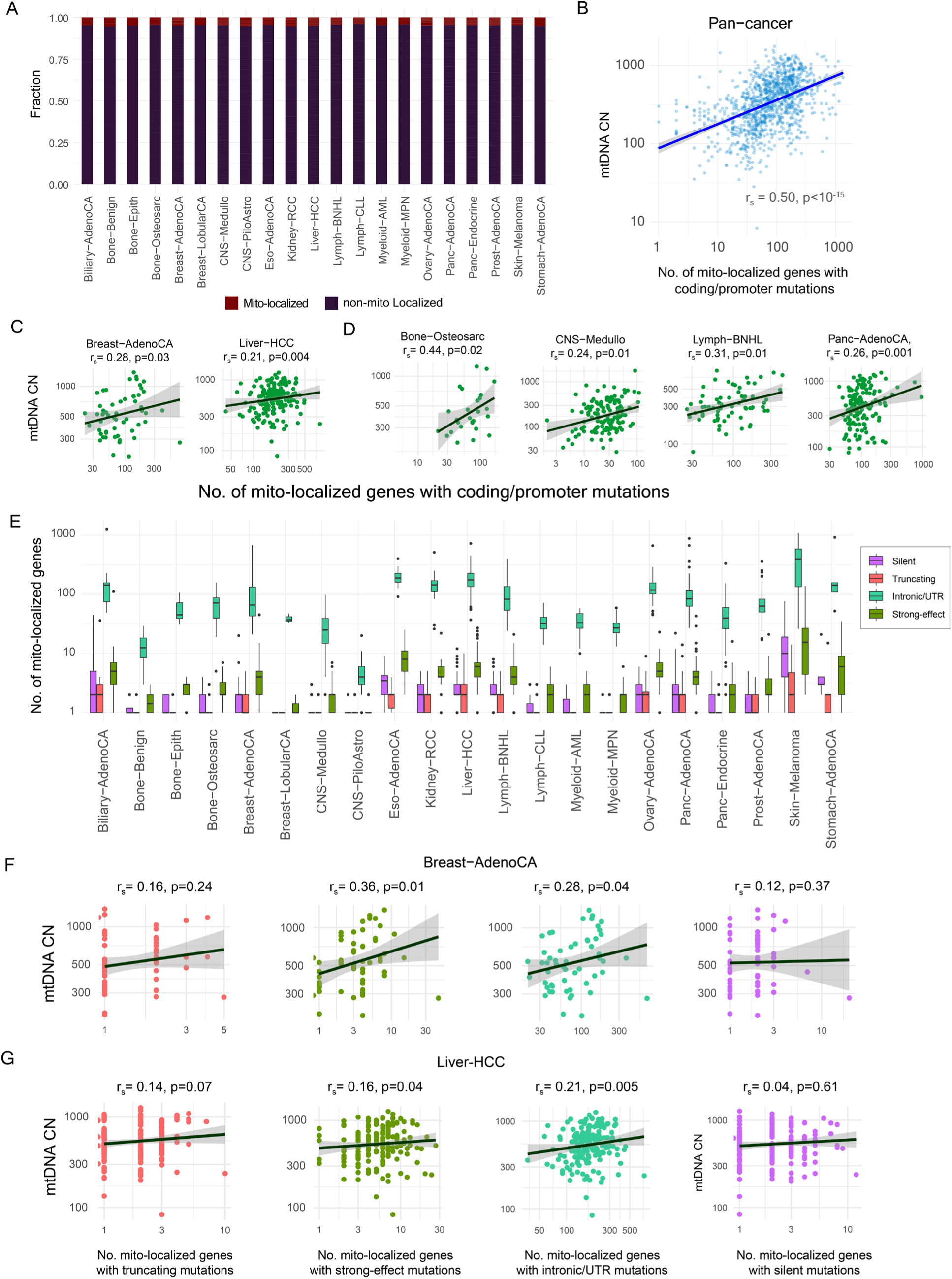
Changes in mtDNA copy number are correlated with the number of mutation-affected genes associated with mitochondrial function. **(A)** Average fraction of mutation-affected nuclear genes that localize to mitochondria in individual cancer types. **(B)** Pan-cancer correlation between the number of mitochondria-localized genes that carry coding and/or promoter mutations and mtDNA copy number. **(C-D)** Correlation between the number of mitochondria-localized genes with coding/promoter mutations and mtDNA copy number in individual cancer types. (C) shows correlations for cancer types which did not originally show any correlation between mutational load and mtDNA copy number, whereas (D) shows correlations for cancer types which originally showed correlation. **(E)** Boxplot showing the distribution of the number of mitochondria-localized genes carrying silent, truncating, intronic/UTR and strong-effect mutations. **(F-G)** Correlation of the number of mitochondria-localized genes with truncating, strong-effect, intronic/UTR and silent mutations with mtDNA copy number in breast adenocarcinoma (F) and liver-HCC (G). The solid lines in scatter plots show linear fits to the data and the shaded grey regions show 95% confidence intervals.

Mutations can affect gene functions to different extent and in different ways. Thus, we further characterized mutations occurring in coding regions of mitochondria-localized genes across individual cancer types (Fig. 3E). Among all mutations observed in coding and promoter regions of mitochondria-localized genes, a large fraction (>50% to ∼90%) were in intron, 3’UTR and 5’UTR regions in all cancer types (Fig. 3E and Fig. S12). These were likely to affect transcription and splicing. Only a small fraction of the mutations was either truncating in nature (frameshift indels and non-sense mutations), or likely to have strong effects on gene function (non-synonymous substitutions and in-frame indels) (Fig. 3E and Fig. S12).

The association between mtDNA copy number and the number of mutations varied according to different categories of mutation and was also cancer type specific. We did not observe any correlation between mtDNA copy number and the number genes affected silent mutations in any of the cancer types (Fig. S13, S14), as silent mutations would have very little fitness effect which did not require functional compensation. However, for strong-effect mutations and intronic/UTR mutations we observed significant correlations between the number of mitochondria-localized genes with such mutations and mtDNA copy number for some of the cancer types (Fig. 3F,G and Fig. S13). For breast adenocarcinoma and liver HCC, we observed significant positive correlation of mtDNA copy number with the number of mitochondria-localized genes with strong-effect and intronic/UTR mutations in (Fig. 3F,G), even though originally there was no correlation between mtDNA copy number and the overall mutational load in these cancer types (Fig. 1G and Fig. S5). Overall, these results suggested that the changes in mtDNA copy number were likely compensating for the loss or reduction in mitochondrial function caused by truncating, strong-effect, or intronic/UTR mutations.

Next, we asked whether the increased mutational load had any role in driving the changes in mtDNA copy number in cancer samples. To address this question, we obtained a list of 49 genes that have been linked to regulation of mtDNA copy number in human (Gupta et al., 2023) and checked whether one or more of these genes were affected by mutations in cancer samples. Across all cancer types, several of these genes contained mutations in coding and/or promoter regions (Fig. S15), suggesting that some of the mutations accumulated in the cancer samples could possibly drive the changes observed in mtDNA copy number in cancer samples.

### Cancer-specific correlation of mtDNA copy number with mutational signatures and structural variants

Next, we tested whether the changes in mtDNA copy number were associated with specific type of nucleotide substitutions or signatures of specific mutagenesis process. We observed variation in specific type of substitutions in cancer-specific manner. For, example, the percentage of A to C and T to G substitutions were markedly higher in esophagus adenocarcinoma compared to other cancer types (Fig. S16A). Although there was weak or no pan-cancer correlation between mtDNA copy number and the number of specific nucleotide substitutions (Fig. S16B), we observed significant correlations for some specific mutational signatures in specific cancer types (Fig. S16C). Similarly, APOBEC signature enrichment had very weak pan-cancer correlation with mtDNA copy number (Fig. S17). However, specific cancer types, such as prostate adenocarcinoma and pancreatic adenocarcinoma, showed significant positive correlation with some of the APOBEC signatures (Fig. S17C).

Pan-cancer analysis of structural variants (SVs) revealed significant positive correlation of the mtDNA copy number with the total number of structural variants (r_s_=0.47, p<10-15; Fig. 4A) as well as with the number of specific types of SVs (Fig. S18). Intriguingly, two cancer types – breast adenocarcinoma and kidney-RCC, which did not originally show correlation between mtDNA copy number and mutational load – had significant correlations between mtDNA copy number and the number of SVs (Fig. S19). We also observed significant variation in occurrence of specific type of SVs in different cancer types (Fig. 4B) and their correlation with mtDNA copy number (Fig. 4C-E). Taken together, these results suggested that the changes in mtDNA copy number is also linked to the accumulation of structural variants in cancer genomes and not just SNVs and small indels.

**Figure 4.**
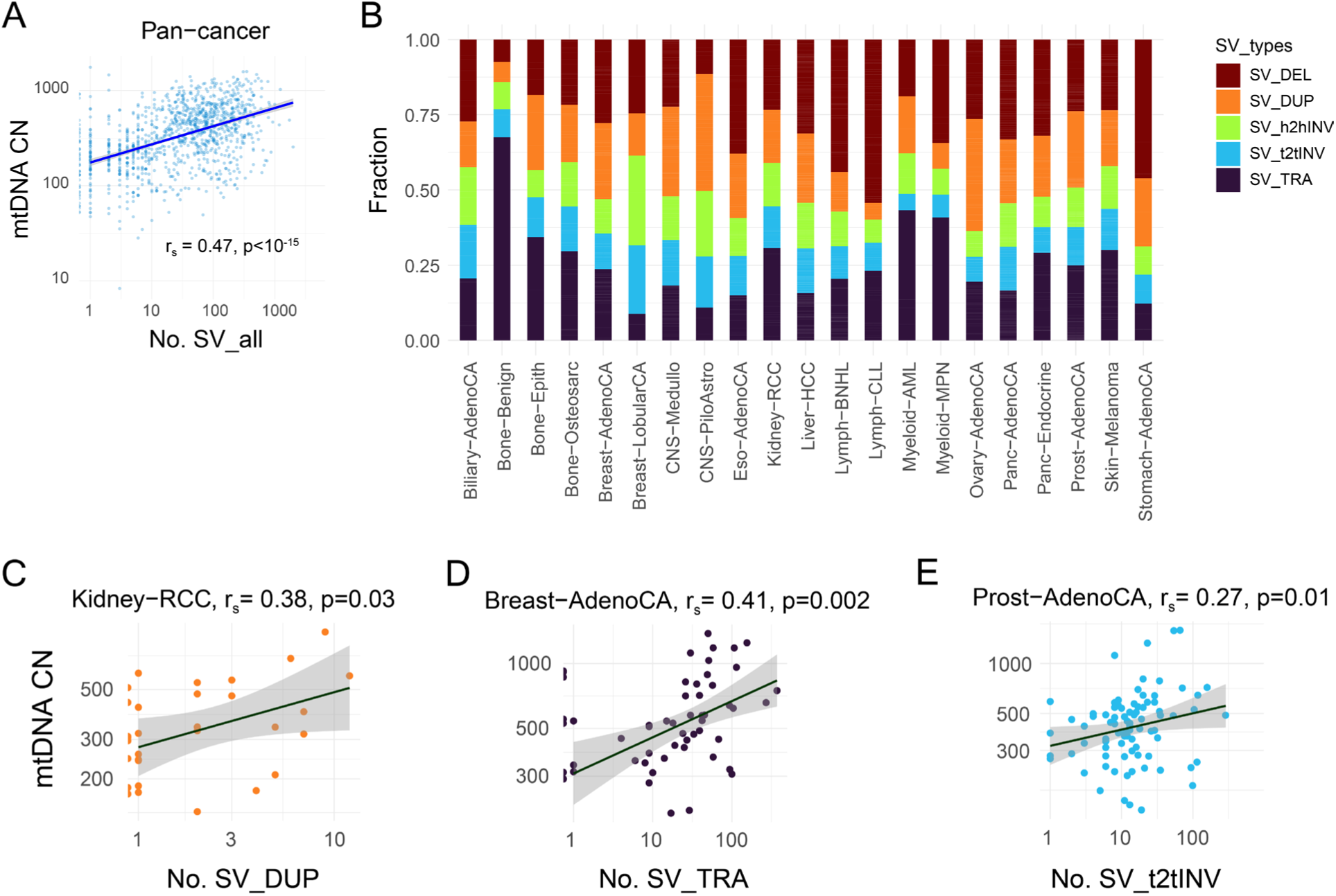
MtDNA copy number is correlated with the occurrence of structural variants in cancer-specific manner. **(A)** Pan-cancer correlation between the number structural variants (of all types) and mtDNA copy number. **(B)** Average fraction of different types of structural variants in individual cancer types. **(C-E)** Correlation of mtDNA copy number with the number of duplications in kidney-RCC (C), translocations in breast adenocarcinoma (D) and tail-to-tail inversions in prostate adenocarcinoma (E). The solid lines in scatter plots show linear fits to the data and the shaded grey regions show 95% confidence intervals.

### Variation in mtDNA copy number leads to differential expression of genes among samples within individual cancer types

To further understand how mtDNA copy number variation can influence cellular processes and phenotypes of cancer cells, we obtained gene expression data for 26 cancer types and considered samples with tumor purity of at least 0.5 for further analyses (Table S4). For each cancer type, we divided samples into low- and high-copy groups based on median mtDNA copy number (Fig. S20). Out of the 26 cancer types, 23 cancer types had at least 3 samples for each of the low- and high-copy number groups (Table S4) and were considered for differential expression analysis.

Next, we performed differential gene expression analyses between the high and low mtDNA copy number groups across cancer types using DESeq2 and identified the differentially expressed genes (FDR<10%, Benjamini-Hochberg correction) (Fig. 5A). The number of upregulated and downregulated genes varied between cancer types with the CNS-GBM showing the lowest and the Liver-HCC showing the highest number of differentially expressed genes respectively (Table S5 and Fig. 5B). For several cancer types, the high- and low-mtDNA samples showed a degree of separation in their expression profiles in principal component analysis (Fig. 5C and Fig. S21). For differentially expressed genes, patterns of change in expression between low- and high-mtDNA copy number samples were visible, despite the widespread genetic differences between samples within individual cancer types (Fig 5D and Fig. S22).

**Figure 5.**
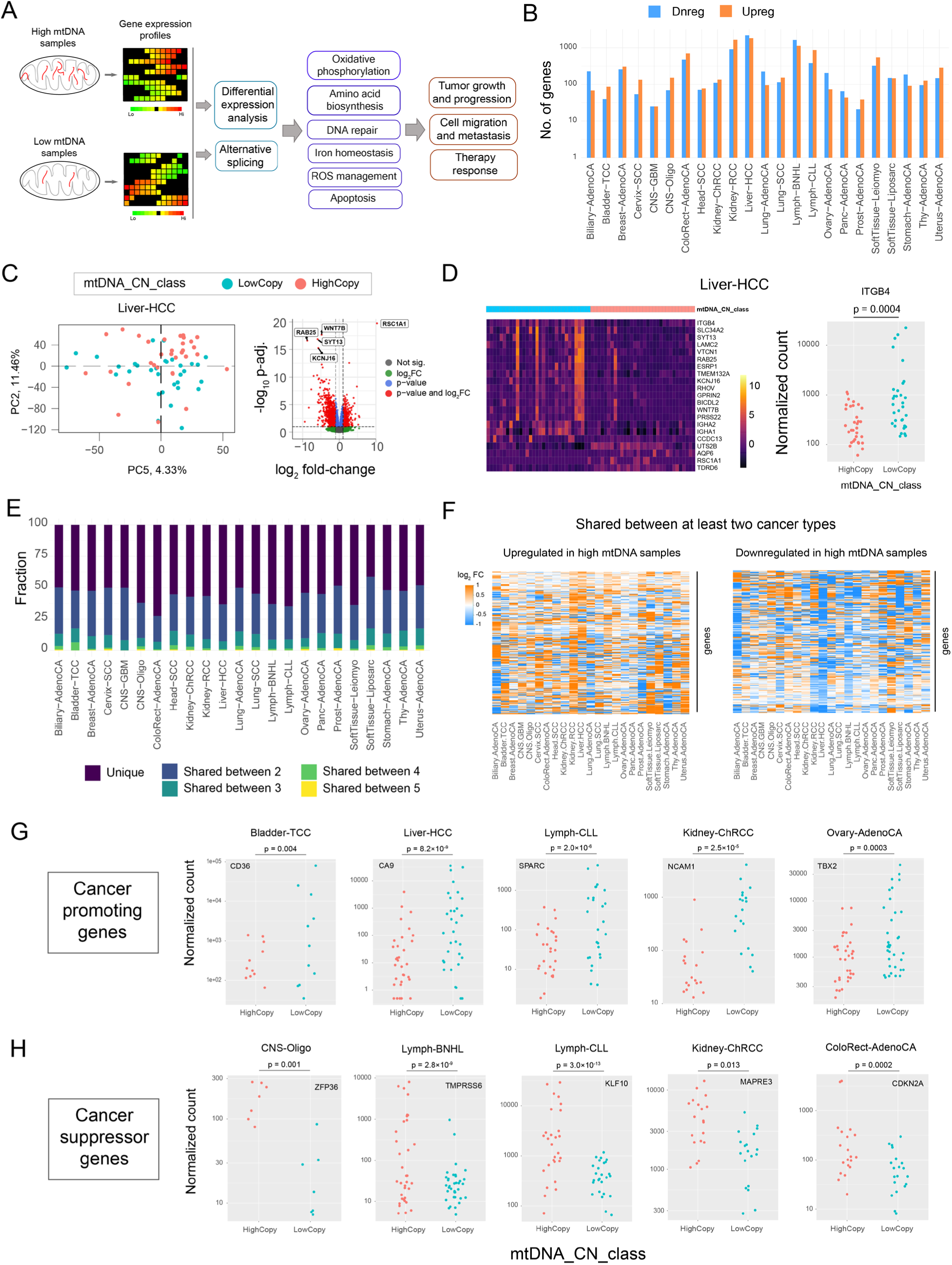
Variation in mtDNA copy number drives differential expression of genes among samples within individual cancer types. **(A)** Schematic diagram showing analysis of differential gene expression and alternative splicing between high- and low-mtDNA samples in individual cancer types, followed by subsequent analysis of the influence of differential expression on cellular pathways and cancer phenotypes. **(B)** Number of genes up- and down-regulated in differential expression analysis across cancer types. **(C)** PCA analysis of high-and low-mtDNA samples in liver-HCC (left), and volcano plot showing the log_2_ fold change and -log_10_p-adjusted values of all genes obtained from differential expression analysis in the same cancer type (right). **(D)** Heatmap showing expression (normalized count) of top 40 genes (identified by differential expression analysis) in low- and high-mtDNA samples in liver-HCC (left) and expression of the gene *ITGB4* in low- and high-mtDNA samples in liver-HCC (right). **(E)** Fraction of unique and shared differentially expressed genes across all cancer types. **(F)** Log_2_ Fold change value of up- and down-regulated genes in high mtDNA samples and shared between at least two cancer types. **(G-H)** Expression of a few representative cancer promoting genes (G) and cancer suppressor genes (H) in high- and low-mtDNA samples in different cancer types.

Among the upregulated genes in high mtDNA copy number groups, 1570 genes were shared between 2 or more cancer types and 36 were shared between more than 4 cancer types (Table S6 and Fig. 5E). Among the down-regulated genes in high mtDNA copy number group, 1455 genes were shared between 2 or more cancer types and 30 were shared between more than 4 cancer types (Table S6 and Fig. 5E). Log fold change of up- and down-regulated genes shared between cancer types (three or more) revealed more than 2-fold change in expression of several genes across different cancer types (Fig. 5F).

### Low mtDNA samples show increased expression of genes and pathways linked to cancer progression and metastasis

Many of the top genes identified in differential expression analysis were associated with cancer progression, and metastasis. Interestingly, several of these genes were upregulated in low mtDNA samples across different cancer types (Fig. 5G and Fig. S23). For example, the gene *CA9*, that showed increased expression in low mtDNA samples in colorectal adenocarcinoma (Fig. 5G), has been shown to promote cancer cell migration and metastasis (Shin et al., 2011). The gene *TBX2*, a known oncogene (Bellis et al., 2025), was upregulated in low mtDNA samples in ovarian adenocarcinoma (Fig. 5G).

On the other hand, several top genes that were upregulated in high mtDNA samples were associated with cancer suppression (Fig. 5H and Fig. S24). For example, the gene *TMPRSS6*, that showed high expression in high mtDNA samples in Lymph-BNHL (Fig. 5H), has recently been found to inhibit neuronal tumor growth (Zuo et al., 2024). Similarly, the gene *DHRS2*, that was upregulated in high mtDNA samples in Head-SCC (Fig. S24), has been shown to inhibit cancer growth and metastasis in ovarian cancer (Li et al., 2022). However, there were exceptions and some of the cancer promoting genes were found to be upregulated in high mtDNA samples of some cancer types (Fig. S25).

For a more comprehensive picture of the cellular transformation of low- and high-mtDNA samples, we performed a functional enrichment analysis of the differentially expressed genes that were observed in at least two cancer types. The genes that were upregulated in high mtDNA samples of at least two cancer types were significantly enriched (FDR<5%) for transmembrane transporter function, including for metal ions, response to low oxygen and detoxification of reactive oxygen species (Fig. 6A). In addition, these genes were also enriched for functions that are likely to support tumor growth such as glutamate biosynthesis, amino acid biosynthesis, carboxylic acid metabolism, and circulatory system processes (Fig. 6A).

**Figure 6.**
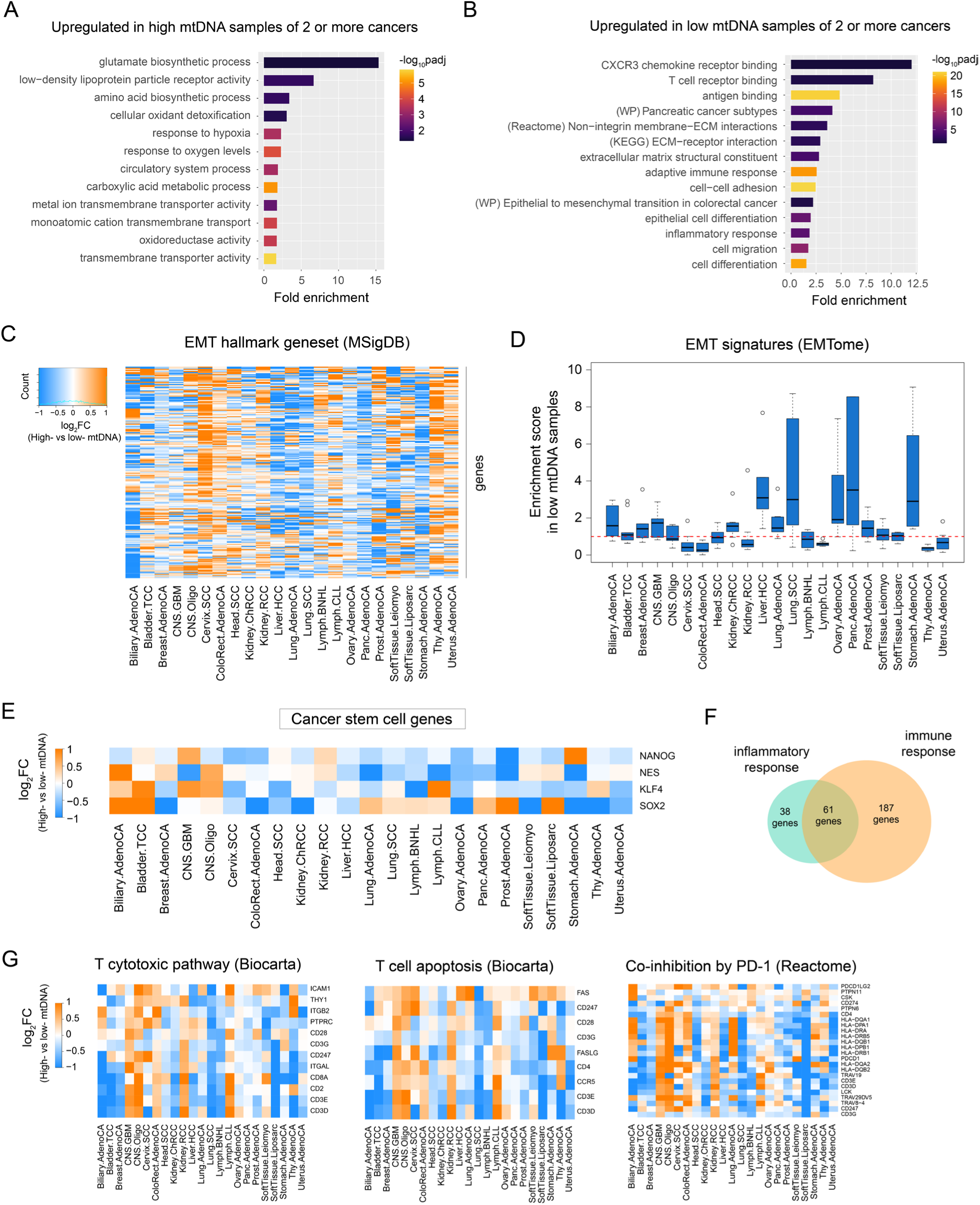
Low mtDNA samples show upregulation of cell migration and EMT genes. **(A)** Functional categories upregulated in high mtDNA samples of two or more cancer types. **(B)** Functional categories upregulated in low mtDNA samples of two or more cancer types. **(C)** Log fold change values (high-vs low-mtDNA) of genes in the EMT hallmark gene set (MSigDB) in different cancer types. **(D)** Enrichment scores in low mtDNA samples calculated for pan-cancer signature EMT gene sets from EMTome database (Vasaikar et al., 2021) across different cancer types. **(E)** Log fold change values (high-vs low-mtDNA) of four genes associated with cancer stemness. **(F)** Venn diagram showing the number of shared and unique genes associated with immune and inflammatory response among genes upregulated in low mtDNA samples of two or more cancer types. **(G)** Log fold change (high-vs low-mtDNA) of genes associated with T cytotoxic pathway (Biocarta), T cell apoptosis (Biocarta) and Co-inhibition by PD-1 pathway (Reactome).

The genes that were upregulated in low mtDNA samples compared to high mtDNA samples in at least two cancer types were significantly enriched (FDR<5%) for cell migration, epithelial cell differentiation, and epithelial to mesenchymal transition (EMT) (Fig. 6B). In addition, we also observed significant enrichment of immune response, T cell receptor binding, and inflammatory response functions (Fig. 6B).

We further investigated for differential expression of EMT genes in low mtDNA samples. To do so, we checked the log fold change (high-vs low-mtDNA) in expression of EMT hallmark genes from the MSigDB database (Fig. 6C) (Liberzon et al., 2015) and several core EMT signature gene sets from the EMTome database (Vasaikar et al., 2021). From EMTome signature gene sets, we also calculated an enrichment score to test whether EMT genes were preferentially upregulated in low mtDNA samples (Fig. 6D). We observed an enrichment of upregulated EMT signature genes in low mtDNA samples (Enrichment score > 1) in 11 cancer types (Fig. 6D). The enrichment scores were particularly high for Liver-HCC, Lung-SCC, pancreatic adenocarcinoma and stomach adenocarcinoma (Fig. 6D). Further, we also observed increased expression of genes associated with cancer stemness such as *NANOG, NES, KLF4 and SOX2* in low mtDNA samples across several cancers (Fig. 6E). We also observed higher expression of cancer stemness genes in cancer specific manner - such as *CD36* in Bladder-TCC and *PROM1* in lung adenocarcinoma.

Out of the 248 genes associated with immune response that showed differential expression between high- and low-mtDNA samples, ∼25% genes were also part of inflammatory response (Fig. 6F). In addition, analysis of signature gene sets revealed upregulation of genes associated with T cell cytotoxicity in low mtDNA samples (Fig. 6G). However, at the same time, the low mtDNA samples also showed signatures of upregulation of genes associated with T cell apoptosis as well as co-inhibition by PD-1 (Fig. 6G).

### Expression of mtDNA genes can predict expression of metabolic and p53 pathways

To further investigate how variation in mtDNA copy number can drive expression of cellular pathways that are relevant for cancer, we built a predictive linear regression model. As expected, mtDNA copy number could well predict the expression of mtDNA encoded genes (Fig. S26). We used the expression level of a representative mtDNA encoded gene, *MT-CO2*, as a marker for the mitochondrial functional state and used its expression as the predictor variable in the linear regression model.

We used the regression model to predict expression of genes in MSigDB hallmark gene sets. As part of the predictive modeling approach, we divided the expression data of each gene into training (80%) and test (20%) data. We fit a linear regression model on the training data, applied the fitted model for predicting expression on the test data, and subsequently calculated Spearman’s correlation coefficient between the predicted expression values and the actual expression values for each gene. We then calculated a mean correlation value for every hallmark gene set by considering predictions for every gene in the gene set.

Among the hallmark pathways, the gene set associated with oxidative phosphorylation showed the highest mean correlation between predicted and actual expression values and with the highest percentage of genes showing significant correlation (p<0.05) (Fig. 7A). Interestingly, the top pathways where a substantial percentage of genes showed significant correlation between predicted and actual expression values included p53 pathway, DNA repair, reactive oxygen species (ROS) response, unfolded protein response and mTORC1 signaling pathways (Fig. 7A,B). The top pathways also included metabolic pathways such as fatty acid metabolism and heme metabolism (Fig. 7A,B). A substantial fraction of genes in each pathway had significant expression correlation with the expression of *MT-CO2* gene (Fig. 7B,C and Fig. S27).

**Figure 7.**
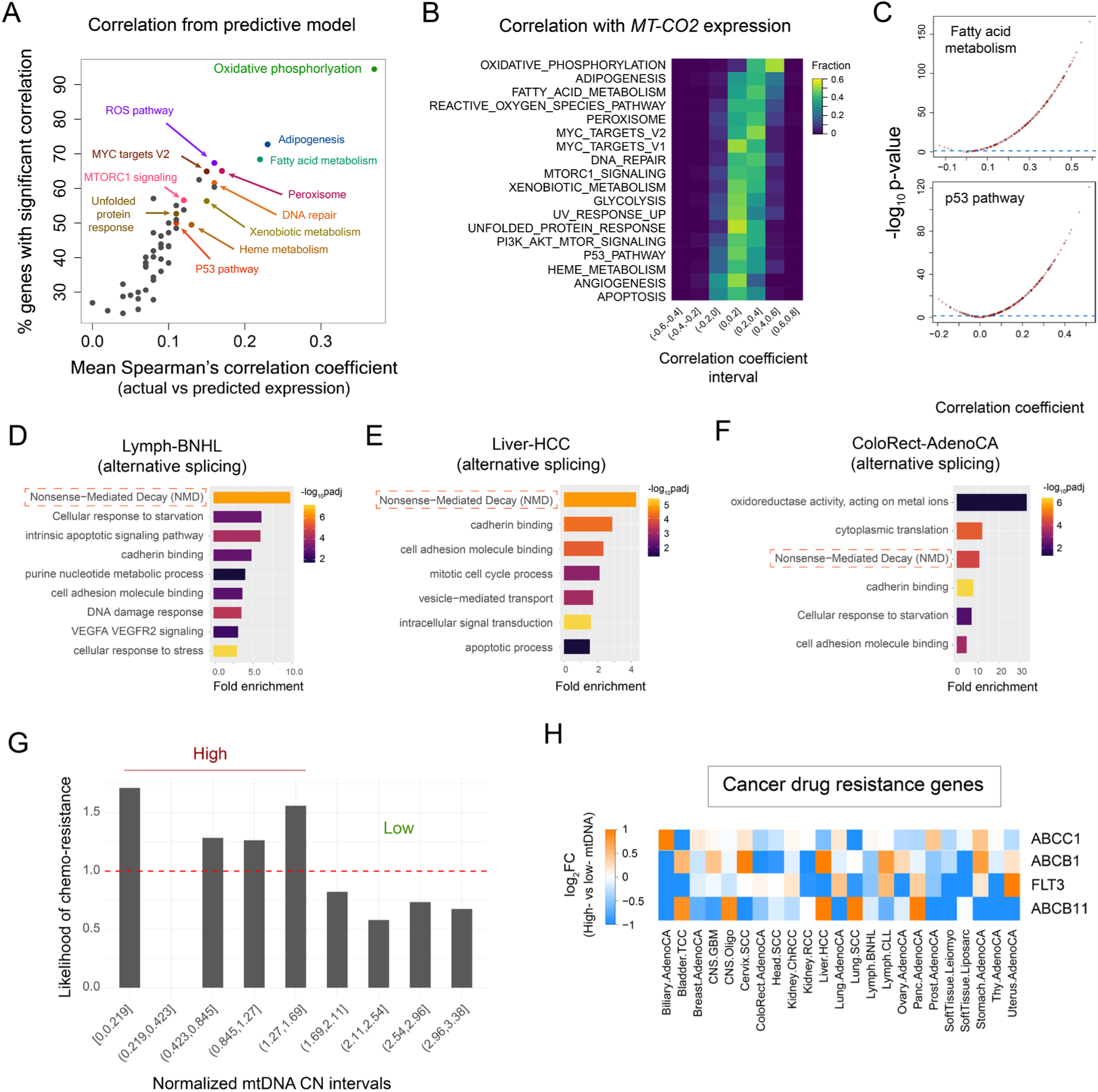
Mitochondrial functional state can predict expression of genes associated with p53 pathway, DNA repair, ROS pathway, and metabolism. **(A)** Scatterplot showing the mean Spearman’s correlation coefficient between actual expression and predicted expression obtained from the regression model for hallmark gene sets and the percentage of genes in the gene sets showing significant correlation (p<0.05). **(B)** Heatmap showing fraction of genes in hallmark gene sets with specific ranges of Spearman’s correlation coefficient values calculated between expression of a gene and MT-CO2 expression. **(C)** Distribution of Spearman’s correlation coefficient and -log_10_(p-value) of correlations calculated between mtDNA encoded MT-CO2 gene and the genes in the hallmark gene sets associated with fatty acid metabolism (top) and the p53 pathway (bottom). **(D-F)** Enriched functions among genes showing alternative splicing between low-vs high-mtDNA samples in Lymph-BNHL (D), Liver-HCC (E) and ColoRect-AdenoCA (F). **(G)** Likelihood of chemotherapy resistance calculated for samples grouped according to normalized mtDNA copy number. **(H)** Log fold change in expression (high-vs low-mtDNA) of three ABC transporter genes (*ABCC1, ABCB1* and *ABCB11*) and *FLT3*.

### Variation in mtDNA copy number is linked to alternative splicing of NMD genes

As a large fraction of mutations in cancer samples were observed in intronic and UTR regions across cancer types (Fig. 3E), we tested for alternative splicing between high-and low-mtDNA copy number samples across cancer types. To do so, we performed differential expression of transcripts between high- and low-mtDNA copy number groups and analyzed the changes in transcript expression (FDR<10%; Benjamini-Hochberg correction). Genes where we observed simultaneous upregulation of at least one transcript and downregulation of at least one transcript were alternatively spliced. Thirteen cancer types showed at least one alternatively spliced gene, with colorectal adenocarcinoma, liver-hepatocellular carcinoma and lymph-BNHL showing many alternatively spliced genes (Table S7). For example, the genes *CD74, RACK1, CDKN1A* and *CIRBP* – all of which have been linked to cancer progression or suppression (Cazier et al., 2014; Liu et al., 2016; Dan et al., 2020; Chen et al., 2021) – showed alternative splicing in Liver-HCC, colorectal adenocarcinoma and Lymph-BNHL, respectively (Fig. S28).

Functional analysis of alternatively spliced genes in three cancer types with substantial numbers of such genes (colorectal adenocarcinoma, liver-hepatocellular carcinoma, and lymph-BNHL) revealed significant enrichment for genes associated with non-sense mediated decay (NMD) and cadherin binding in all these cancer types (Fig. 7D-F). In addition, we observed enrichment of genes associated with cellular response to starvation, DNA damage response, apoptotic process and signaling for apoptotic process in one or more of these cancer types (Fig. 7D-F).

### Low mtDNA copy number increases the likelihood of chemotherapy resistance

Variation in mtDNA copy number significantly altered expression of genes linked to chemotherapy response such as *TBX2, CES1, DUSP1*, and *ITGB4* (Shen et al., 2016; Lin and Zhu, 2024; Jiang et al., 2024; Bellis et al., 2025). Thus, we tested whether the likelihood of developing chemotherapy resistance had any association with mtDNA copy number. To do so, we considered samples with known chemotherapy treatment and known therapy outcomes across all cancer types. To address the issue of inherent variation in mtDNA copy number among different cancer types, we used a measure of normalized mtDNA copy number derived through median normalization across cancer types. At low values of normalized mtDNA copy number, we observed higher likelihood of developing chemo-resistance (likelihood > 1), whereas the likelihood dropped below 1 at increased levels of normalized mtDNA copy number (Fig. 7G). These results suggested that low mtDNA copy number is likely to increase the chances of developing chemoresistance in cancer.

Further, we tested for expression of genes associated with drug transport and resistance. Although the genes *ABCB1* and *ABCC1*, that are known to be involved in drug efflux in cancer (Xiao et al., 2021; Engle and Kumar, 2022; Bergonzini et al., 2024) showed upregulation in low mtDNA samples of only some cancer types, another transporter *ABCB11* (Hayashi and Sugiyama, 2013) showed higher expression in low mtDNA samples in several cancers (Fig. 7H). Similarly, another gene *FLT3*, that has been linked to resistance to FLT3 inhibitors (Ruglioni et al., 2024) also showed higher expression in low mtDNA samples across several cancer types (Fig. 7H).

## Discussion

Taken together, our results reveal a central role of mitochondria in cancer progression. We observed an overall positive correlation between mtDNA copy number and mutational load in cancer samples. The mtDNA copy number could also predict the extent of mutational load, suggesting a causal link between the two in cancer. In addition, the tumor ploidy, and the number of nuclear indels were the top predictors of the mtDNA copy number. These results highlighted likely causal links between these features and mtDNA copy number and hinted at a possible compensatory role of increased mtDNA copy number in cancer samples.

The signature of functional compensation by increased mtDNA copy number was further strengthened from the analysis of functions of genes that were affected by mutations. In individual cancer types, mtDNA copy number was positively correlated with the number of mitochondria-localized genes that carried either truncating, strong-effect, or intronic/UTR mutations, suggesting possible compensation for the detrimental effects caused by mutations in genes linked to mitochondrial functions.

In several cancer samples, the mtDNA copy number increased with an increase in tumor ploidy but without a concomitant increase in mutational load (Fig. S10). This suggested that changes in mtDNA copy number perhaps occurred first which could then allow further accumulation of mutations without a substantial detrimental effect on mitochondrial function. However, in prostate adenocarcinoma, we observed an increase in mutational load without an increase in mtDNA copy number (Fig. S11), suggesting that there could also be cancer-specific sequence of events. In addition, we also observed a positive correlation between the number of structural variants and the mtDNA copy number in several cancer types where we did not see any correlation between mtDNA copy number and SNVs/indels. These results suggested that in cancer samples, an increase in mtDNA copy number could allow not just accumulation of SNVs/indels, but also structural variants for which functional compensation might be necessary.

Variation in mtDNA copy number led to differential expression of many genes in cancer samples. High mtDNA samples showed increased expression of pathways that were likely to enable tumor growth and survival in the tumor microenvironment compared to the low mtDNA samples. On the other hand, low mtDNA samples often showed increased expression of genes associated with cell migration and epithelial-to-mesenchymal (EMT) transition (Fig. 6B-D). Interestingly, expression level of a mtDNA encoded gene, *MT-CO2*, could predict the expression of the p53 pathway (Fig. 7A,B), suggesting that mitochondrial activity could be an essential component for maintaining this important tumor suppressor pathway in cancer. In addition, mitochondrial functional state could predict expression levels of DNA repair and unfolded protein response pathways as well as several metabolic functions (Fig. 7A), further highlighting how mitochondrial functional state could regulate cancer progression.

Further, our work linked variation in mtDNA copy number to alternative splicing in genes associated with non-sense mediated decay (NMD). NMD is a quality control mechanism that primarily degrades transcripts with premature stop codons and regulates gene expression (Popp and Maquat, 2013; Nogueira et al., 2021). Although NMD can facilitate cancer progression by suppressing expression of tumor suppressors and enabling immune evasion (Mort et al., 2008; Lindeboom et al., 2016), it is unclear how alternative splicing of NMD genes can affect cancer progression. It has been suggested that the efficiency of NMD can rise with increased mutational burden in cancer (Zhao and Pritchard, 2019). Thus, it is possible that alternative splicing of the NMD associated genes in fact boosts the efficacy of the NMD machinery.

Taken together, these findings reveal how variation in mtDNA copy number and corresponding changes in mitochondrial functional state can influence cancer progression. Although high mtDNA samples accumulate a substantially higher number of mutations compared to low mtDNA samples, the detrimental effects of these mutations are likely to be compensated by increased mtDNA copy number. Further, high mtDNA samples show upregulation of tumor suppressors and the p53 pathway genes. In contrast, the low mtDNA samples show upregulation of genes linked to cell migration, differentiation, and epithelial-mesenchymal transition, as well as increase the likelihood of chemoresistance. Therefore, these findings establish a central role for mitochondria and its functional state in determining mutational load, cancer progression, and therapy response. Thus, altering mitochondrial activity through designed interventions could lead to more effective disease management and anti-cancer therapy.

## Methods

### Dataset

The mutation data were obtained from the PCAWG project (PCAWG 2020), and the public part of the data were downloaded from the ICGC data portal (Zhang et al., 2019). Only the high-quality consensus SNVs, MNVs, and small INDELs that passed quality filtering in the PCAWG project were used in our analyses. In addition, data on consensus SVs, consensus CNVs, and mutational signatures including APOBEC mutagenesis data were analyzed. The tumor purity data for all samples were obtained from the consensus CNV call data. The data on first therapy type and first therapy response were obtained from clinical and histology data of the PCAWG project.

The data on mtDNA copy number of tumor samples were obtained from The Cancer Mitochondria Atlas (TCMA) project (Yuan et al., 2020). In addition, the data on SNVs and indels in the mtDNA were also obtained from the TCMA project. To compare mtDNA copy numbers of the tumor samples with the normal samples of the same cancer type, the aligned whole-genome and whole-exome read data from normal tissue, normal blood, or other normal samples (e.g., bone marrow, lymph node) as available for each cancer type, were analyzed. Briefly, the aligned read data was processed to calculate genome-wide read coverage in normal samples using *bedtools* (Quinlan and Hall, 2010). Subsequently, the mtDNA copy number was calculated from the ratio of the mtDNA coverage to the nuclear DNA coverage.

### Mutation mapping and functional classification

The SNVs/Indels were classified into promoter, coding, intronic/UTR, and intergenic mutations. Mutations that were in the 5’ flanking region of a gene and within 1000 bp upstream of the start codon were considered as promoter mutations. For promoter mapping, the GRCh37 assembly version of the human genome was used. The coding mutations were further classified as truncating, strong-effect, and silent mutations. The truncating mutations consisted of frameshift indels and nonsense mutations, whereas the strong-effect mutations consisted of in-frame indels and missense mutations.

The collated data for patients were further filtered using tumor purity values. Only samples with purity of 0.5 or above were considered for further analyses (Fig. S29). This resulted in a total of 1230 samples of 24 cancer types being included in our analyses (Table S1). Normalized mtDNA copy numbers for individual cancer types were calculated through median normalization and the median mtDNA copy number for each cancer type was set to one.

The nuclear genes that regulate mtDNA copy number in normal human cells were identified by Gupta et al. (2023). A total of 49 such genes were included in our analysis (Table S3). Functional association of genes were done using cellular components (CC), molecular functions (MF), and biological processes (BP) data from Gene Ontology (GO) database (2024 version) (The Gene Ontology Consortium, 2023). The genes that were either localized in mitochondria or were associated with any biological process in mitochondria were identified.

### Correlation analysis of mtDNA copy number with mutational load

Pan-cancer analysis of the correlation of mtDNA copy number with the total mutational load, the numbers of nuclear SNVs, nuclear indels, mtDNA SNVs and mtDNA indels were performed. Spearman’s rank correlation was used for all analyses and the correlation coefficient and the p-values were reported. In addition, pan-cancer analysis of correlation between mtDNA copy number and specific types of nucleotide substitutions and APOBEC enrichment values were done. For pan-cancer analyses, all 1230 samples were included. Subsequently, all correlation analyses were repeated for each cancer type individually. For analysis in individual cancer types, only cancer types with 3 or more samples were included, thereby, resulting in inclusion of 21 cancer types (Table S1).

Further, variation of mtDNA copy number and mutational load with cancer stages were analyzed for all individual cancer types where information was available (Fig. S30 and S31). TNM staging of samples was converted to stages 1 to 4 according to the mapping described by Rosen and Sapra (2023). No consistent trends (increase or decrease in copy number or mutations) with cancer stage were observed across cancer types with a few exceptions.

To test whether mtDNA copy number of a sample can predict its mutational load, the pan-cancer collated data was divided into training (80% of the full dataset) and test (remaining 20% data) data. A linear regression model was fitted to the training data and subsequently, mtDNA copy number of the samples in the test data were used to predict their mutational load. The predicted mutational load and the actual mutational load for the test data were compared using Spearman’s rank correlation.

### Linear regression and random forest models for prediction of mtDNA copy number

To identify the molecular features that can predict mtDNA copy number in cancer samples, two linear regression models and one random forest model were built. The mtDNA copy number was the target variable and a total of 87 features in the collated dataset were used as predictor variables (Table S2). The predictor variables ranged from the number of nuclear and mtDNA SNVs, indels, specific nucleotide substitutions, number of missense, nonsense mutations, APOBEC signature enrichment, to number of genes affected by mutations, number of genes affected by mutations of specific types (truncating/strong-effect/Intron or UTR), and number of mitochondria-localized genes affected by specific types of mutations.

For linear regression models, Lasso and Ridge regression models were used, as many of the features in our data were likely to show significant positive correlation with each other (multi-collinearity). Lasso and Ridge regression can identify the key features for prediction in presence of multi-collinearity and can prevent overfitting. Ridge regression was performed with the package *‘ridge*’ (Cule et al., 2022) and Lasso regression was performed with the *‘glmnet’* package (Friedman et al., 2010) in R. In Ridge regression, the parameters with significant effect on regression (p<0.05) were considered as the key parameters. In Lasso regression, the parameters with non-zero coefficient values were considered as the key parameters. For Lasso regression, the best penalty parameter (lambda) was obtained through a grid search of the parameter value with a 10-fold cross validation. The final regression models were run with only the key parameters and the best lambda value. For the random forest model, the R package ‘randomForest’ (Liaw and Wiener, 2002) with default parameter values was used and the key features were identified through a ranking of features according to their importance.

For prediction, the collated data with only selected key features were taken and the full data were partitioned into training (80% of the full data) and test (remaining 20%) data. The models were first fitted with the training data and subsequently, prediction of mtDNA copy number was done using the test data. The predicted and the actual values were compared with Spearman’s correlation coefficient.

### Differential expression analysis of genes and transcripts between high- and low-mtDNA samples

The transcript data for 26 cancer types (Table S4) were downloaded from the ICGC data portal. The transcript level data were converted to gene level data by taking the sum of all expressed transcripts for a gene. Only samples with tumor purity of 0.5 or more were considered in our analysis. For each cancer type, the median mtDNA copy number was calculated. For a cancer type, the samples with lower mtDNA copy number than the median were classified as low mtDNA samples, whereas the samples with higher mtDNA copy number than the median were classified as high mtDNA samples. Three cancer types (Breast-LobularCA, Cervix-AdenoCA and Eso-AdenoCA) had less than three samples in the low- and high-mtDNA categories and hence, these cancer types were not considered for further analyses.

For each of the remaining 23 cancer types, differential gene expression analysis was performed at both gene and transcript levels using the Bioconductor package *DESeq2* (Love et al., 2014). The differentially expressed genes were identified by a p-adjusted cut-off of 0.1 (FDR<10%, Benjamini-Hochberg method). Principal component analyses of the samples were done using the Bioconductor package ‘*PCAtools’* (Blighe and Lun, 2019). Volcano plots were generated using the package *‘EnhancedVolcano’* (Blighe et al., 2018), and the expression heatmaps were generated using ‘*pheatmap*’ (Kolde, 2025).

Within each cancer type, the upregulated genes in high-mtDNA samples and the upregulated genes in low-mtDNA samples were further checked for their shared up-or down-regulation profile among two or more cancer types. This enabled categorization of unique and common differentially expressed genes across cancer types.

For identification of alternative splicing, the results from the differential expression of transcripts were used. The differentially expressed transcripts were first mapped to their corresponding genes. For a cancer type, genes that showed simultaneous upregulation of one or more transcripts and downregulation of one or more transcripts were considered to be alternatively spliced. Functional enrichment of the alternatively spliced genes was performed using ‘*gprofiler*’ (version *e113_eg59_p19_f6a03c19*, databases updated 2024) (Raudvere et al., 2019), and the functional enrichment for Biological Processes in Gene Ontology (GO) database (database versions: 2024) (GO consortium, 2023).

### Pathway enrichment and analysis of signature gene sets

The functional enrichment analysis for the genes up- and down-regulated in two or more cancer types were performed using ‘*gprofiler*’ (version *e113_eg59_p19_f6a03c19*, databases updated 2024) (Raudvere et al., 2019), and the functional enrichment for Biological Processes in Gene Ontology (GO) database, KEGG pathways (Kanehisa et al., 2014) and Reactome pathways (Milacic et al., 2024) (database versions: 2024) were performed. The significantly enriched classes were obtained by filtering classes with adjusted p-value less than 0.1 (10% False Discovery Rate) and fold enrichment values for these classes were calculated using the following formula.

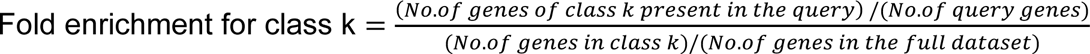

For each cancer type, the fold change in expression of specific signature gene sets associated with specific cellular functions were checked and represented as heatmaps. The signature hallmark gene sets were obtained from MSigDB database and all 50 hallmark gene sets were analysed (Liberzon et al., 2015). In addition, a collection of EMT signature gene sets was obtained from the EMTome database (Vasaikar et al., 2021). The enrichment score of EMT signatures in low mtDNA samples was calculated as follows. From the EMTome database, only the pan-cancer signatures were considered which led to inclusion of 10 signature gene sets. For each signature, the log_2_ fold change and the p-adjusted values of the genes in the signature were obtained. For genes showing significant differential expression (p-adjusted<0.1), the number of genes upregulated in the low mtDNA samples (n_low_) and the number of genes upregulated in the high mtDNA samples (n_high_) were counted. Next, the enrichment score in the low mtDNA samples were calculated as 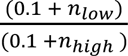. Similarly, signature gene sets related to immune response from reactome and biocarta pathways maps in MSigDB were analysed.

### Regression models for predicting expression of hallmark gene sets

To test whether mitochondrial functional state can predict expression patterns of genes in specific cellular pathways, a predictive model based on linear regression was built. As expected, mtDNA copy number could well predict the expression levels of mtDNA encoded genes (Fig. S26). Among the mtDNA encoded genes, expression levels of one representative gene, *MT-CO2*, was used as an indicator of the mitochondrial functional state.

Next, expression of *MT-CO2* was used to predict the expression levels of genes in hallmark gene sets. For every gene in a hallmark gene set, a linear regression model was set with *MT-CO2* as the predictor variable. The expression data of the target gene and *MT-CO2* were divided into training (80% of the data) and test (remaining 20%) sets. The linear regression models were fitted on the training data and then the fitted model was used for prediction of the target gene expression from *MT-CO2* expression level in the test data. The predicted expression values and the actual expression values of the target gene were compared by Spearman’s rank correlation. For each hallmark gene set, the mean correlation coefficient value, and the number of genes with significant correlation (p<0.05) between actual and predicted values were reported. Subsequently, for the hallmark gene sets, individual correlation values between the expression levels of the genes in the gene set and the expression level of *MT-CO2* were calculated.

### Calculation of likelihood score for chemotherapy resistance

To test whether mtDNA copy number has any influence on response to chemotherapy, a likelihood score for chemotherapy resistance was calculated. Across all cancer types, the patients with chemotherapy as the first therapy and with a known therapy response were included. The therapy response varied from ‘complete response’, ‘stable disease’ and ‘disease progression’ among patients who received chemotherapy. The patients showing ‘disease progression’ after chemotherapy treatment were resistant to chemotherapy, whereas patients showing ‘complete response’ or ‘stable disease’ were responding (fully or partially) to the chemotherapy treatment. To account for the inherent variation in mtDNA copy number among different cancer types, normalized mtDNA copy numbers were calculated by median normalization across cancer types and the median mtDNA copy number were set to one. To calculate the likelihood score for resistance, all included patients were divided into 10 equal sized bins according to their normalized mtDNA copy number. In each bin, the likelihood score was calculated as 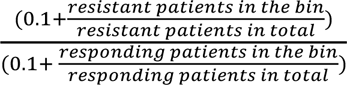. No score was reported for bins without any observation.

## Acknowledgement

We are highly grateful to the International Cancer Genome Consortium (ICGC) for granting access to the PCAWG data. We thank the research participants and their families who donated samples and data. We acknowledge contributions of the physicians and the clinical staff who contributed to sample collection, annotation, and analysis. We also acknowledge contributions of all research groups and clinical networks across ICGC and TCGA, as well as the PCAWG consortium for the collated genomic, transcriptomic, and clinical data from the cancer patients.

## Author contributions

Conceptualization: RD Methodology: AA, RD Investigation: AA, RD Visualization: AA, RD Funding acquisition: RD Project administration: RD Supervision: RD Writing – original draft: AA, RD Writing – review & editing: AA, RD

## Data availability

The genomic and transcriptomic data used in the manuscript are available at the ICGC data portal post data access authorization. The TCMA data are available at https://bioinformatics.mdanderson.org/public-software/tcma/

## Code availability

All codes for data analysis and models are available in github: https://github.com/riddhimandhar/Cancer_mtDNAcn

## Supplementary material

### Supplementary figures

**Figures S1 to S31**

### Supplementary tables

**Table S1** – List of cancer types along with the corresponding number of samples included in mutation analysis

**Table S2** – List of features used for building linear regression and random forest models for predictive modelling of mtDNA copy number

**Table S3** – List of genes linked to mtDNA copy number regulation in human cells obtained from Gupta et al., 2023

**Table S4** – Number of total samples in high- and low-mtDNA copy number groups of 26 cancer types available for gene expression analysis

**Table S5** – Number of up- and down-regulated genes in 23 cancer types obtained from differential expression analysis

**Table S6** – List of up- and down-regulated genes in high mtDNA samples compared to low mtDNA samples and shared between two or more cancer types

**Table S7** - Number of genes exhibiting alternative splicing between high- and low-mtDNA groups

**Table S8** - Genes exhibiting alternative splicing between high- and low-mtDNA groups in ColoRect-AdenoCA, Liver-HCC, Lymph-BNHL

