## Supplementary figures for "Pan-cancer analysis reveals mtDNA copy number as a key determinant of mutational load and disease progression in cancer"

A

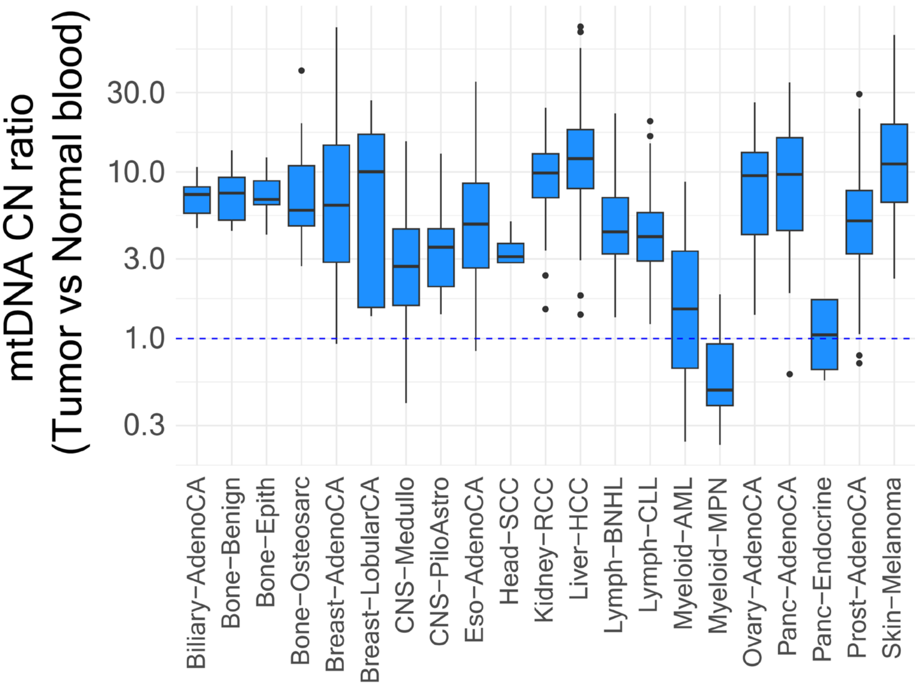


B

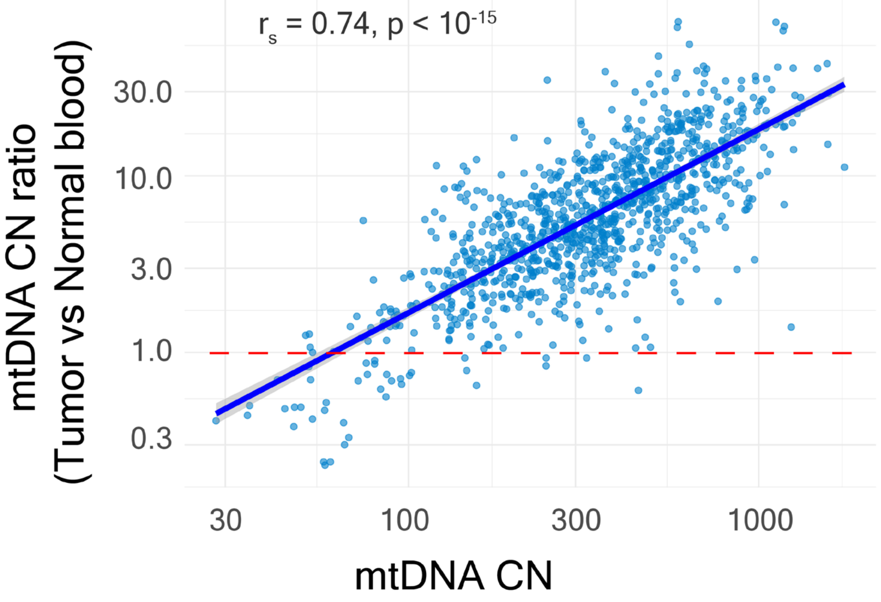


**Figure S1 – (A)** Ratio of mtDNA copy number (mtDNA CN) of the cancer samples to that of the matched normal blood samples. The dotted lines at 1.0 indicates no change in mtDNA copy number in tumor samples compared to normal samples. **(B)** Correlation between mtDNA CN ratio (cancer vs matched normal blood) and mtDNA CN. The solid blue line shows the linear fit to the data and the shaded grey region shows 95% confidence interval.

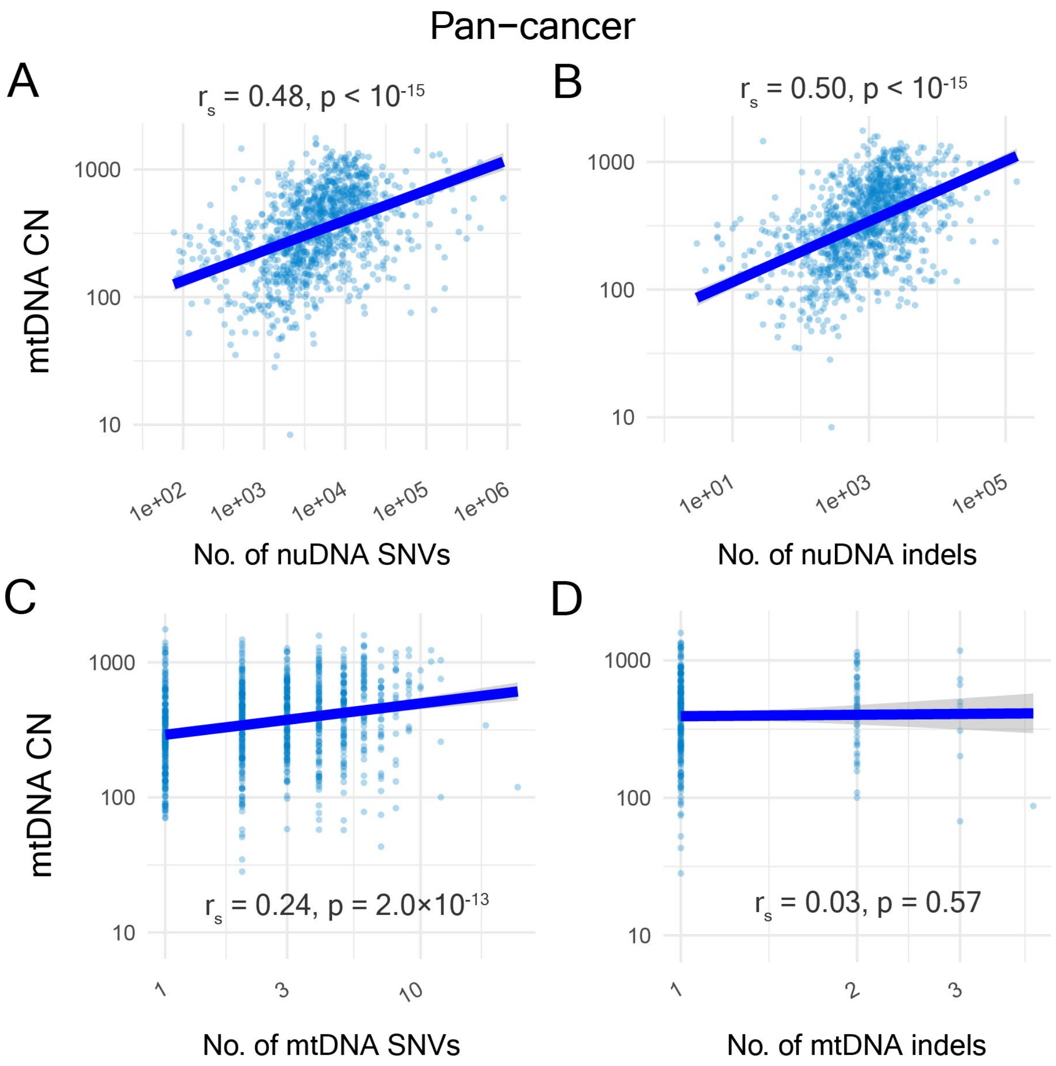


**Figure S2 –** Correlation of mtDNA copy number with **(A)** the number of nuDNA SNVs, **(B)** the number of nuDNA INDELs, **(C)** the number of mtDNA SNVs, and **(D)** the number of mtDNA INDELs. The solid blue lines show linear fits to the data and the shaded grey regions show 95% confidence intervals.

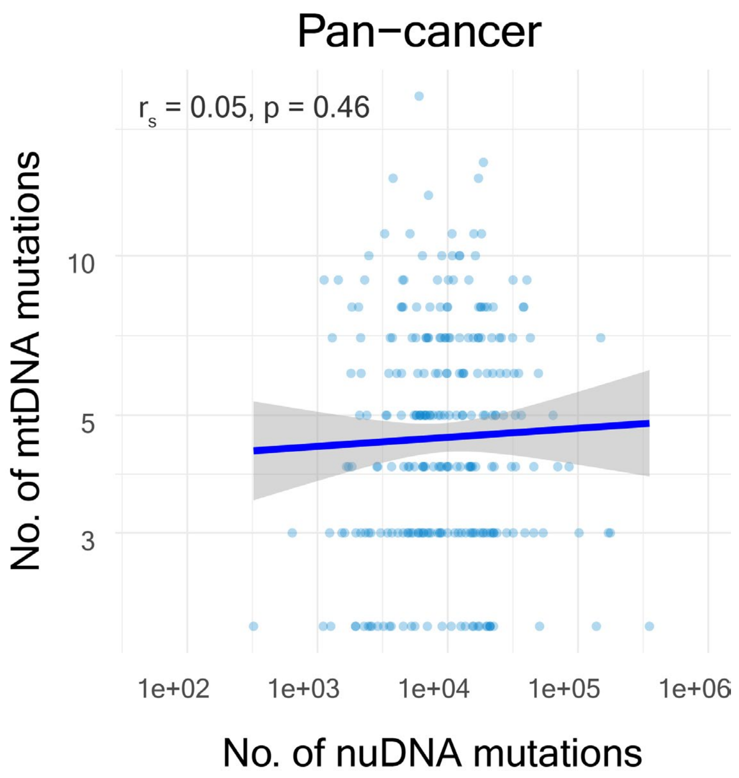


**Figure S3 –** Pan-cancer correlation between the number of nuclear DNA mutations and the number of mtDNA mutations in cancer samples. The solid blue line shows linear fit to the data and the shaded grey region shows 95% confidence interval.

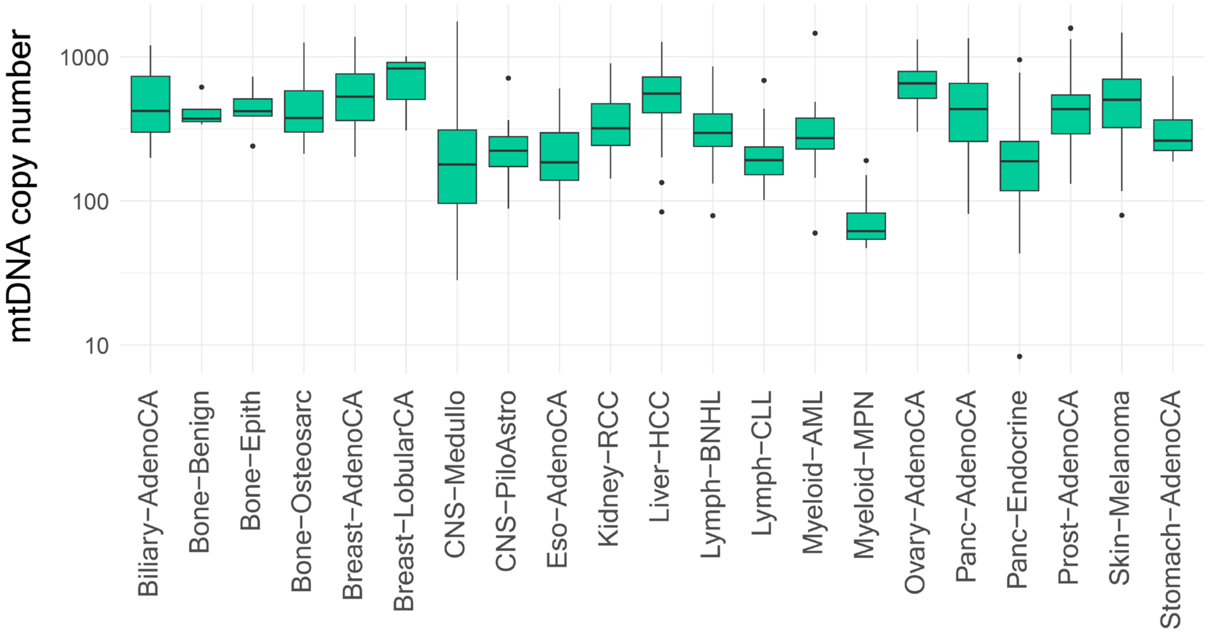


**Figure S4 –** Distribution of mtDNA copy number in individual cancer types included in our analysis.

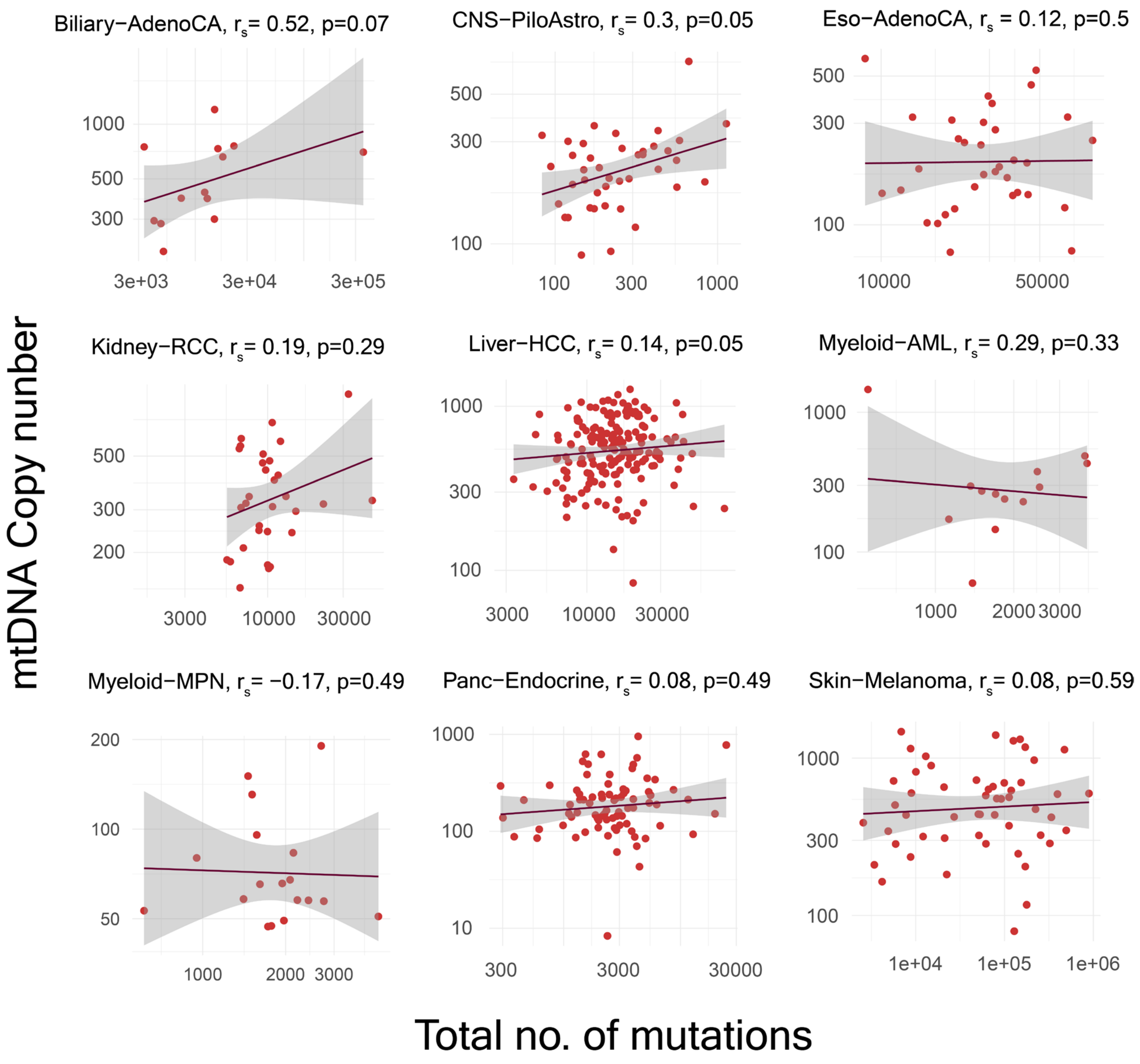


**Figure S5 –** Correlation between mtDNA copy number and the total number of mutations (nuDNA + mtDNA) in individual cancer types. None of these cancer types showed significant correlation between these two features. The solid lines show linear fits to the data and the shaded grey regions show 95% confidence intervals.

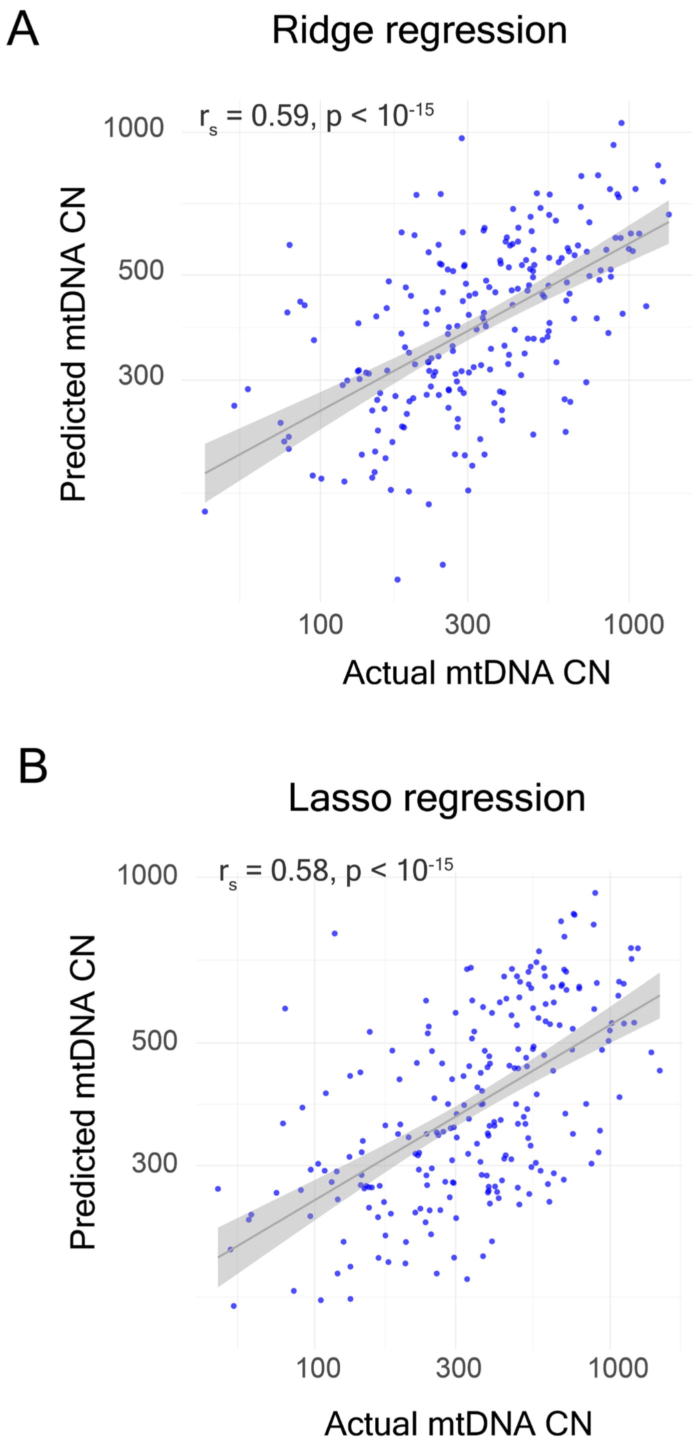


**Figure S6 – (A)** Correlation between predicted mtDNA copy number (CN) by Ridge regression and the actual mtDNA copy number in cancer samples. **(B)** Correlation between predicted mtDNA copy number by Lasso regression and the actual mtDNA copy number in the cancer samples. The solid lines show linear fits to the data and the shaded grey regions show 95% confidence intervals.

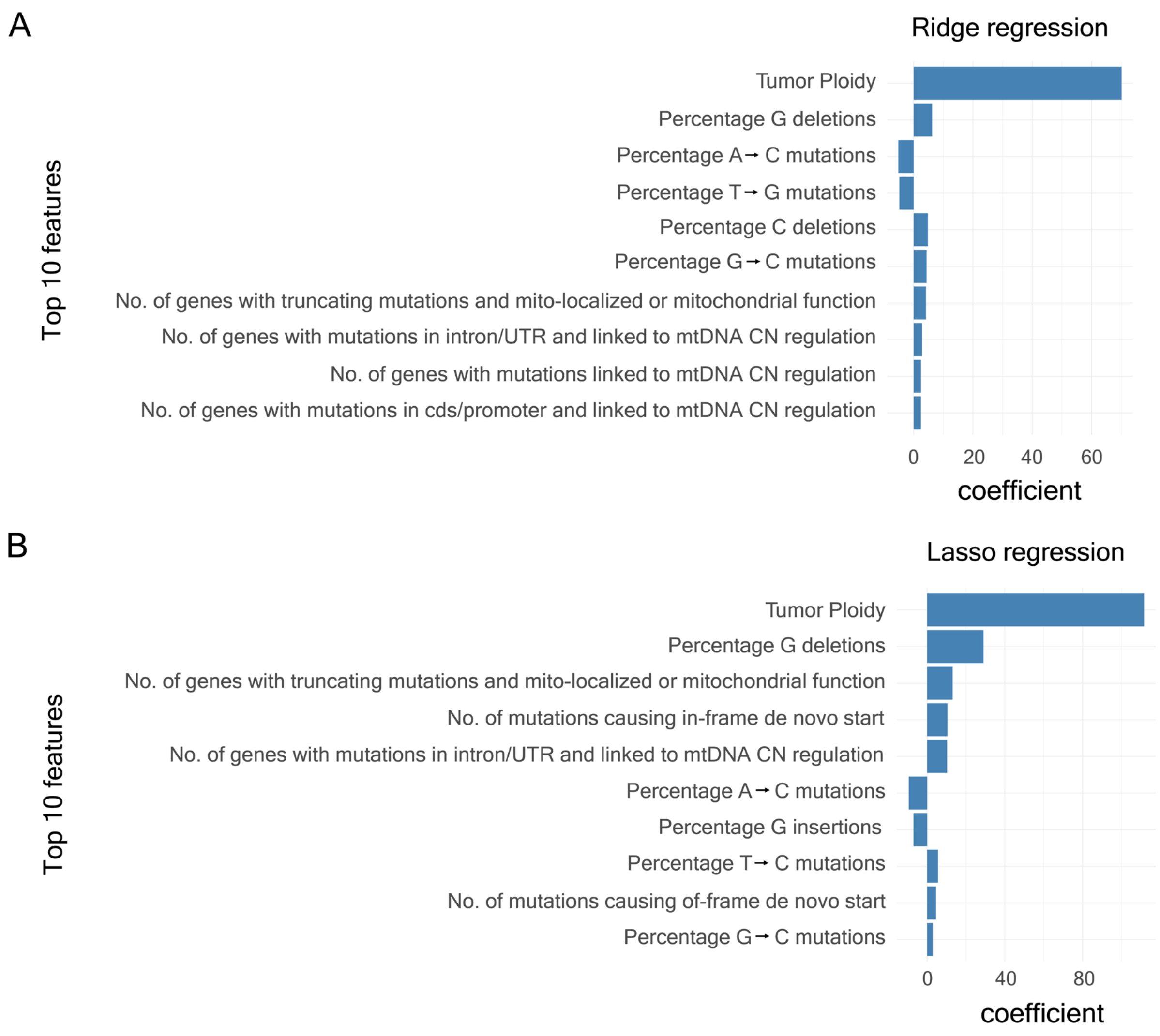


**Figure S7 –** The top 10 features in the predictive regression models ranked by their coefficients in **(A)** Ridge regression and **(B)** Lasso regression.

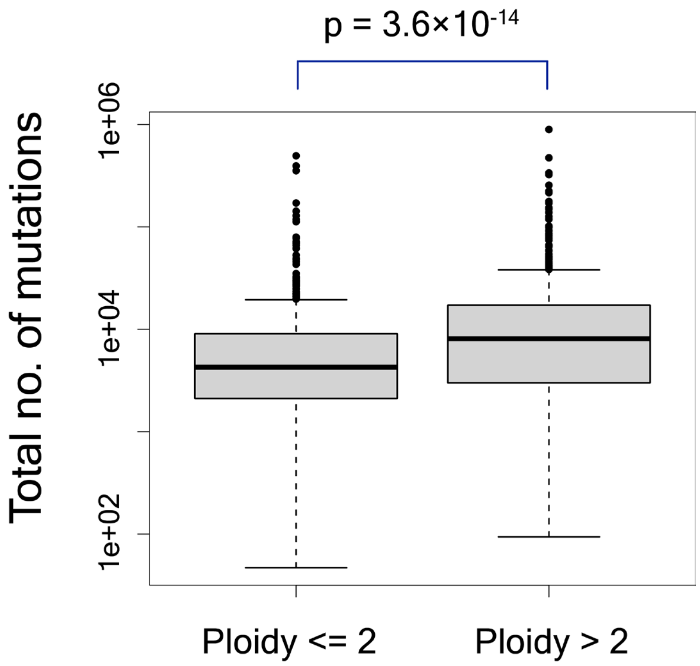


**Figure S8 –** Pan-cancer distribution of the total number of mutations in cancer samples with tumor ploidy ≤ 2 and with ploidy > 2.

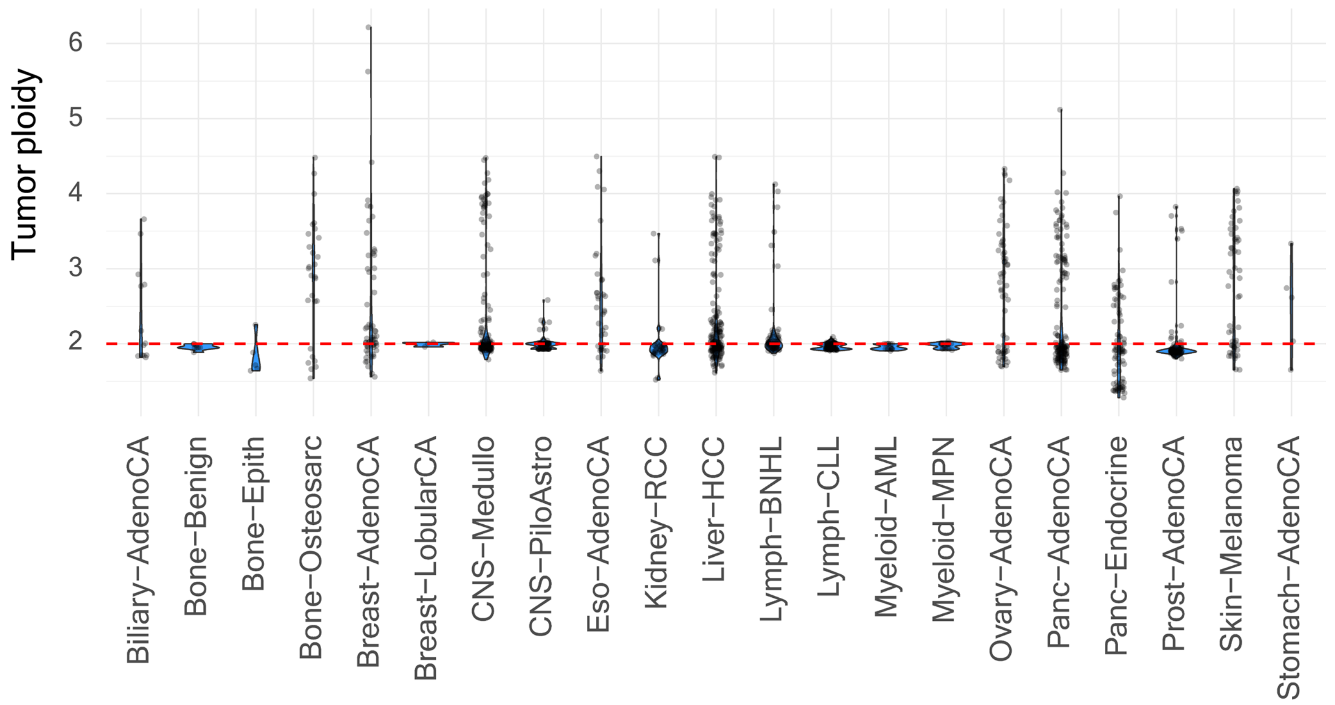


**Figure S9 –** Distribution of tumor ploidy values in individual cancer types. The red dotted line shows the ploidy level 2.

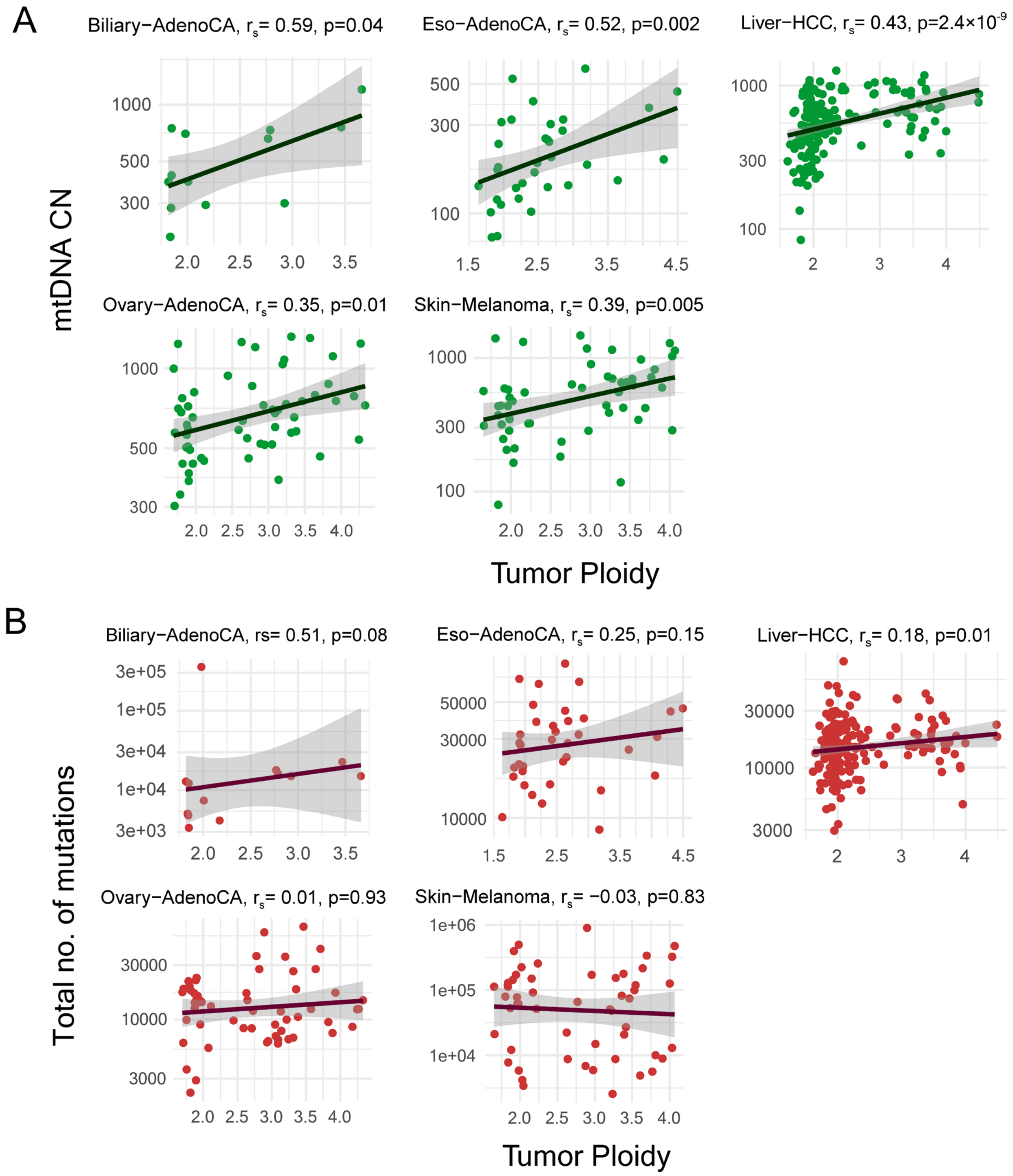


**Figure S10 –** Correlation between Tumor ploidy and **(A)** mtDNA copy number and **(B)** total number of mutations in five cancer types where the mtDNA copy number was significantly correlated with tumor ploidy but the total number of mutations did not show correlation with tumor ploidy except for Liver-HCC. The solid lines show linear fits to the data and the shaded grey regions show 95% confidence intervals.

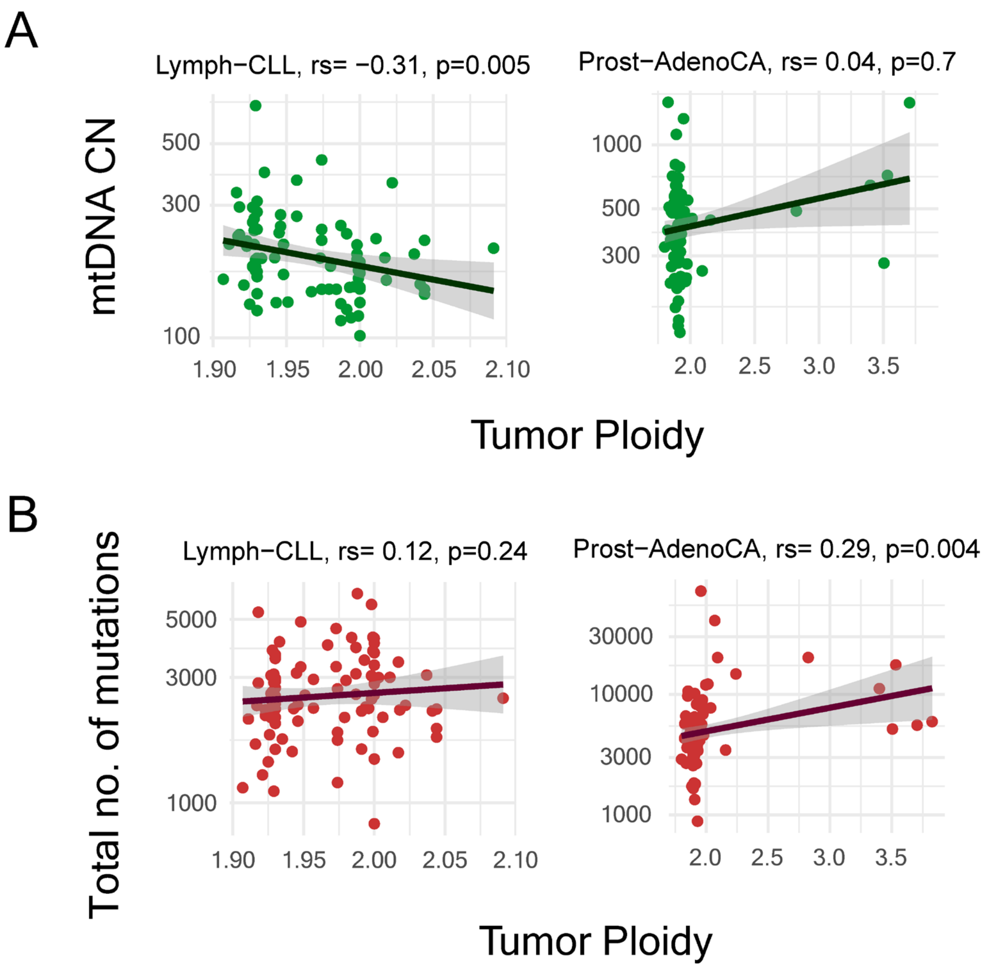


**Figure S11 –** Correlation between Tumor ploidy and **(A)** mtDNA copy number and **(B)** total number of mutations in two cancer types (Lymph-CLL and Prost-AdenoCA). In Lymph CLL, the mtDNA copy number was negative correlated with tumor ploidy, whereas the total number of mutations did not show any correlation with tumor ploidy. In Prostate-AdenoCA, the mtDNA copy number did not show any correlation with tumor ploidy but the total number of mutations was positively correlated with tumor ploidy. The solid lines show linear fits to the data and the shaded grey regions show 95% confidence intervals.

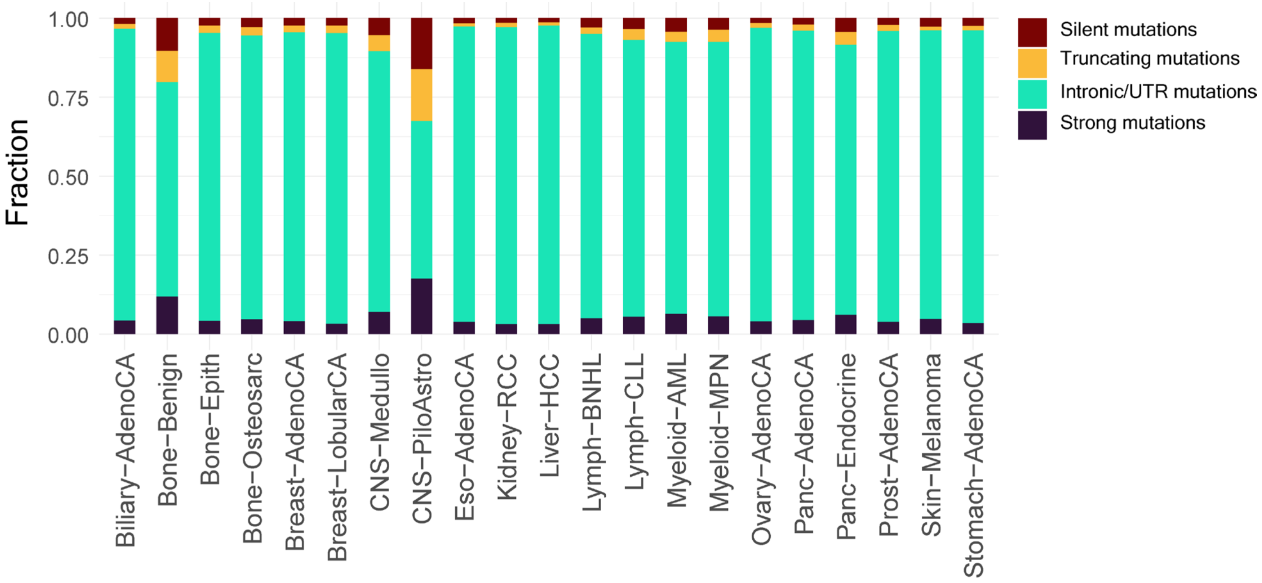


**Figure S12 –** Average fraction of truncating, strong-effect, intronic/UTR and silent mutations in individual cancer types.

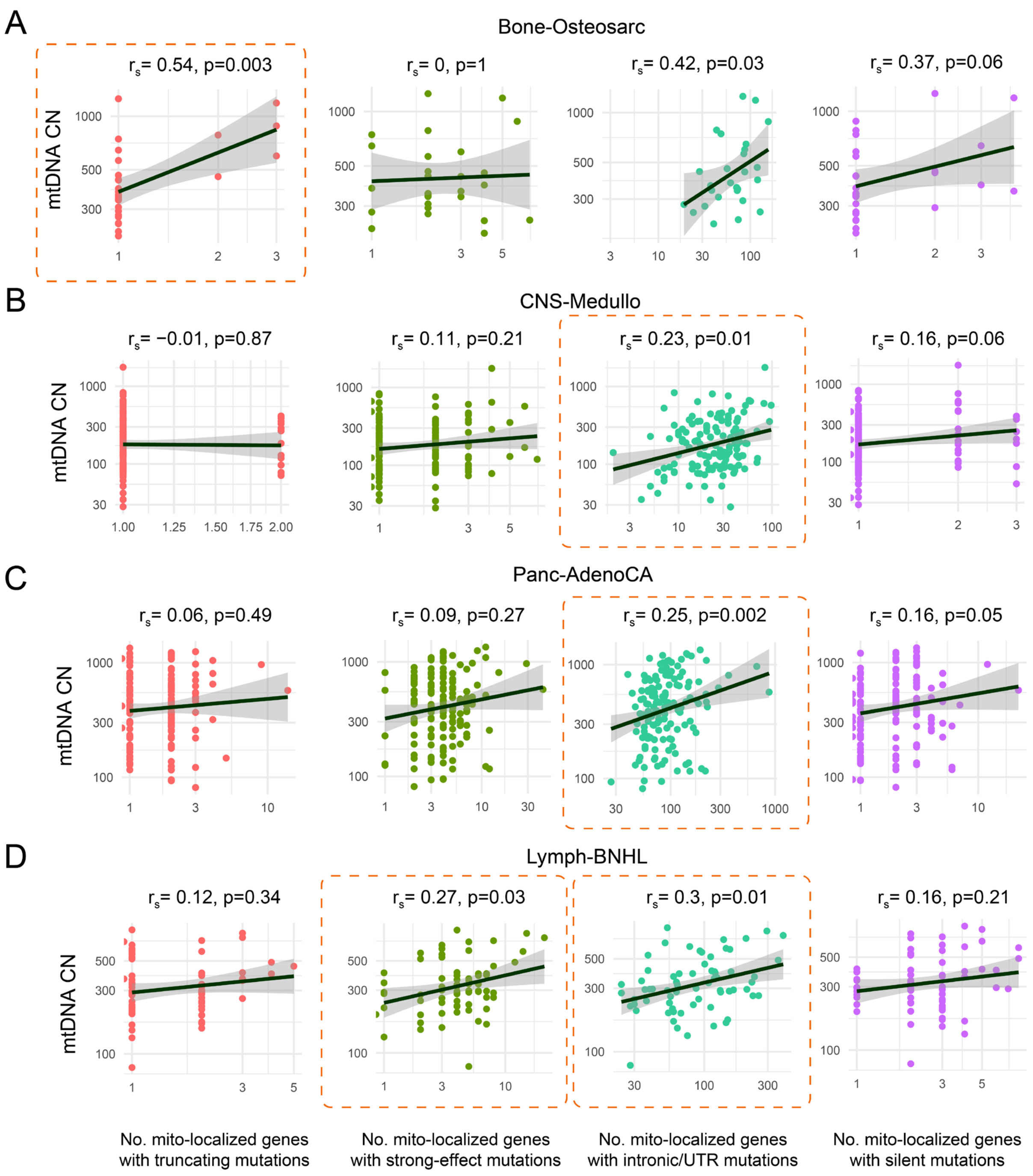


**Figure S13 –** Correlation of the number of mitochondria-localized genes with truncating mutations, the number of strong-effect mutations, the number of intronic/UTR mutations and the number of silent mutations with mtDNA copy number in Bone-osteosarc **(A)**, CNS-Medullo **(B)**, Panc-AdenoCA **(C)**, and Lymph-BNHL **(D)**. In these cancer types, the mtDNA copy number was significantly positively correlated with at least one type of mutation. The solid lines show linear fits to the data and the shaded grey regions show 95% confidence intervals.

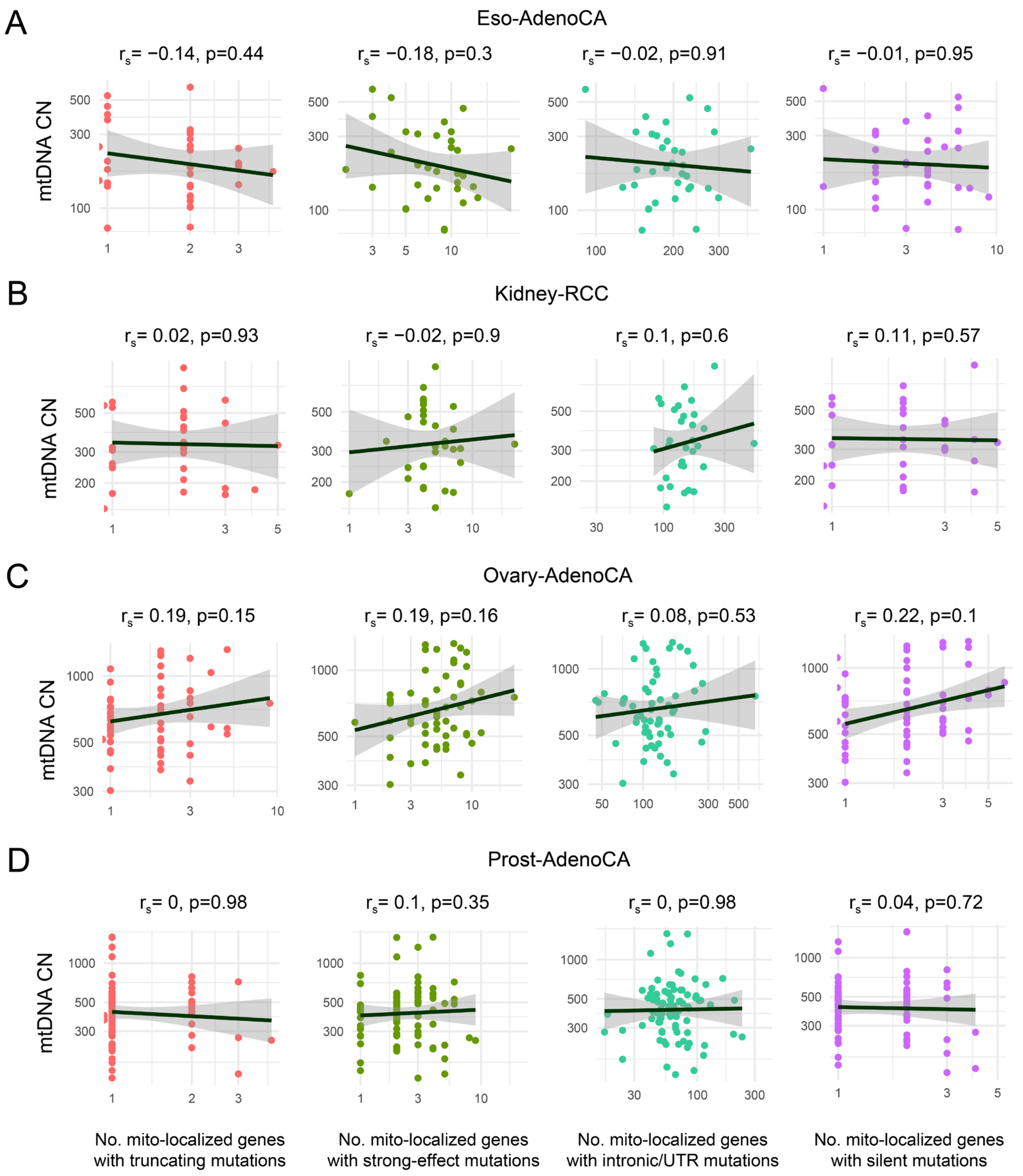


**Figure S14 –** Correlation of the number of mitochondria-localized genes with truncating mutations, strong-effect mutations, intronic/UTR mutations and silent mutations with mtDNA copy number in Eso-AdenoCA **(A)**, Kidney-RCC **(B)**, Ovary-AdenoCA **(C)**, and Prost-AdenoCA **(D)**. In these cancer types, the mtDNA copy number did not show correlation with any of the mutation types. The solid lines show linear fits to the data and the shaded grey regions show 95% confidence intervals.

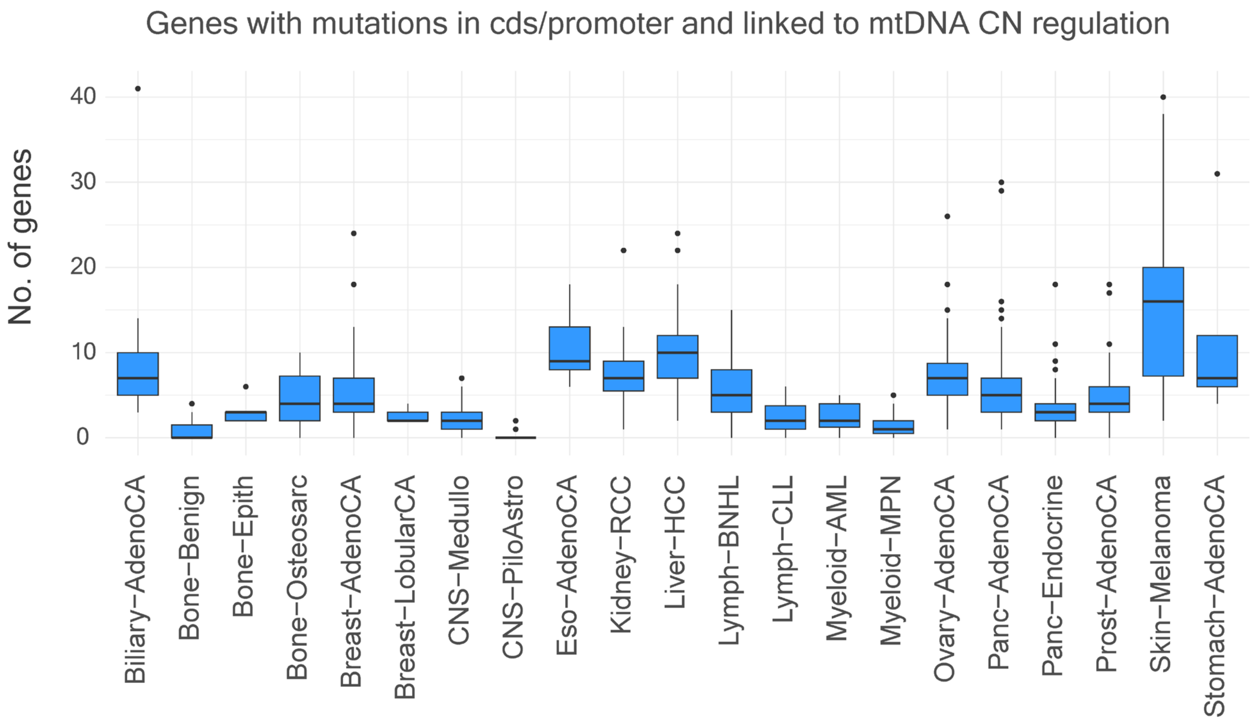


**Figure S15 –** Number of genes affected by mutations in the coding and/or promoter region and have been associated with regulation of mtDNA copy number in human.

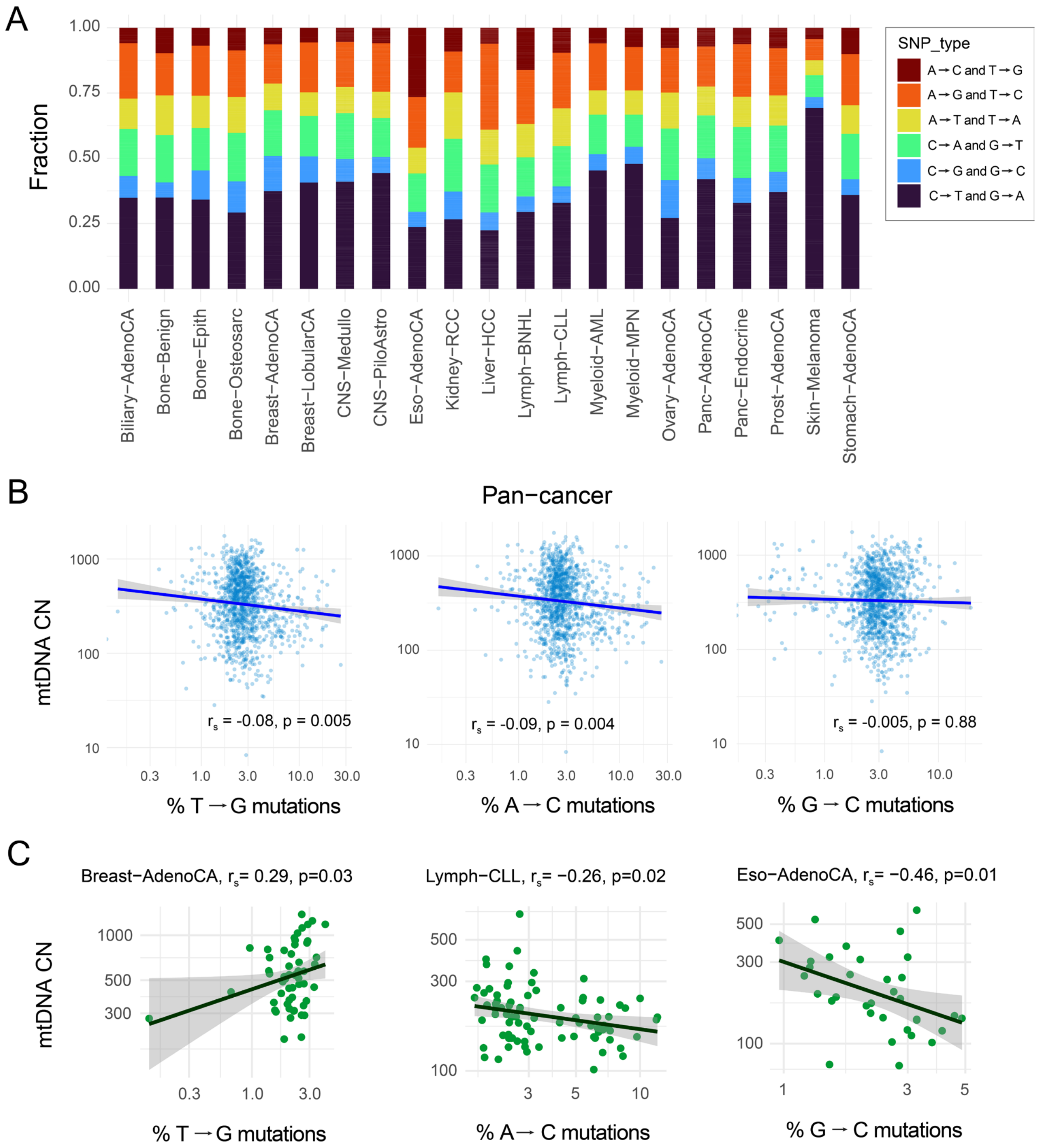


**Figure S16 – (A)** Average fraction of different types of nucleotide substitution observed in individual cancer types. **(B)** Pan-cancer correlation between mtDNA copy number and percentage of T to G, A to C and G to C mutations. **(C)** Correlation between mtDNA copy number and percentage T to G mutations, A to C mutations and G to C mutations in individual cancer types. Only some of the cancer types with significant correlation are shown. The solid lines show linear fits to the data and the shaded grey regions show 95% confidence intervals.

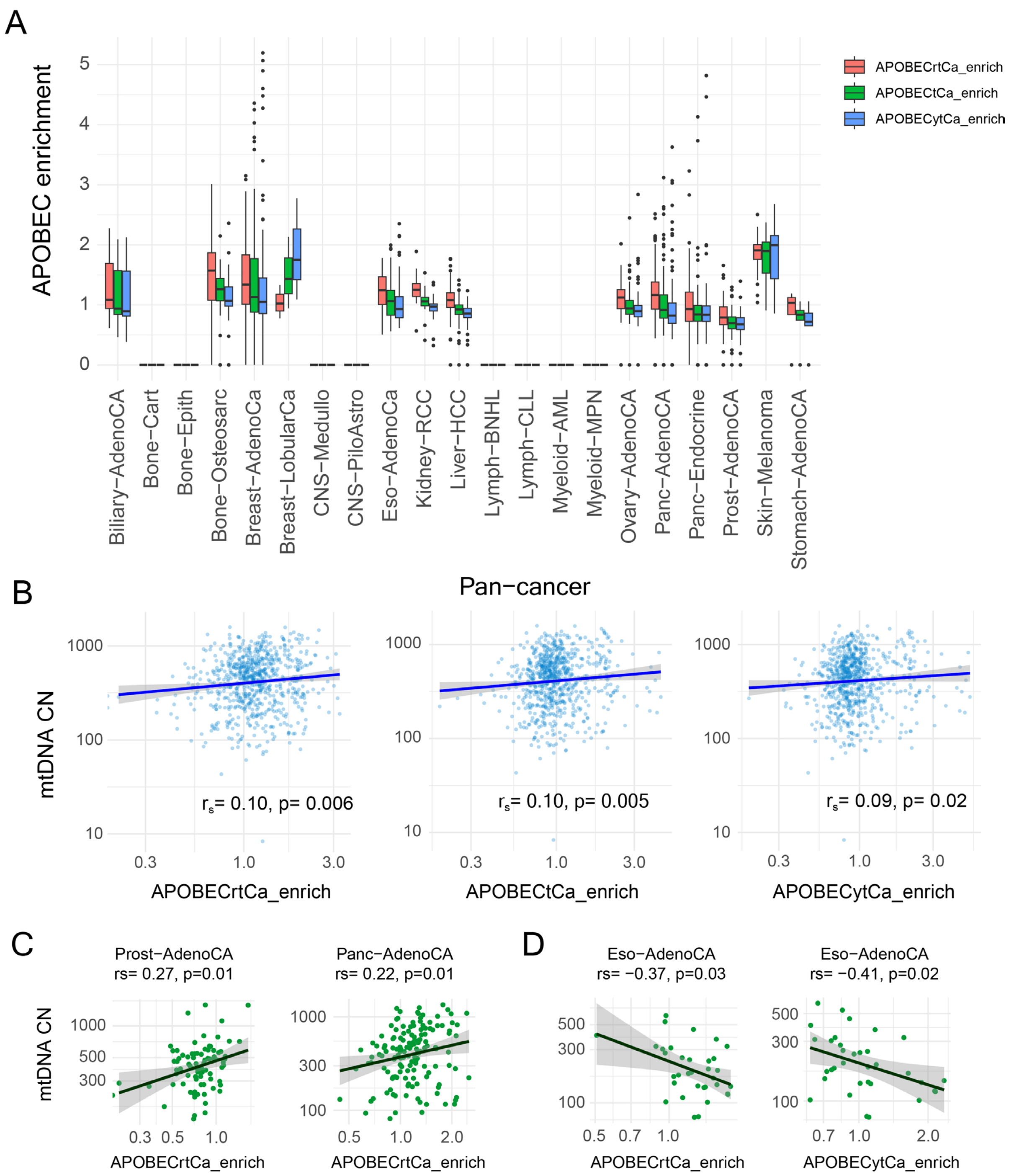


**Figure S17 – (A)** Distribution of APOBEC enrichment scores in individual cancer types. **(B)** Pan-cancer correlation between mtDNA copy number and the APOBEC enrichment scores. **(C)** Correlation between mtDNA copy number and the APOBEC enrichment scores in individual cancer types. Only some of the cancer types with significant correlation are shown. The solid lines show linear fits to the data and the shaded grey regions show 95% confidence intervals.

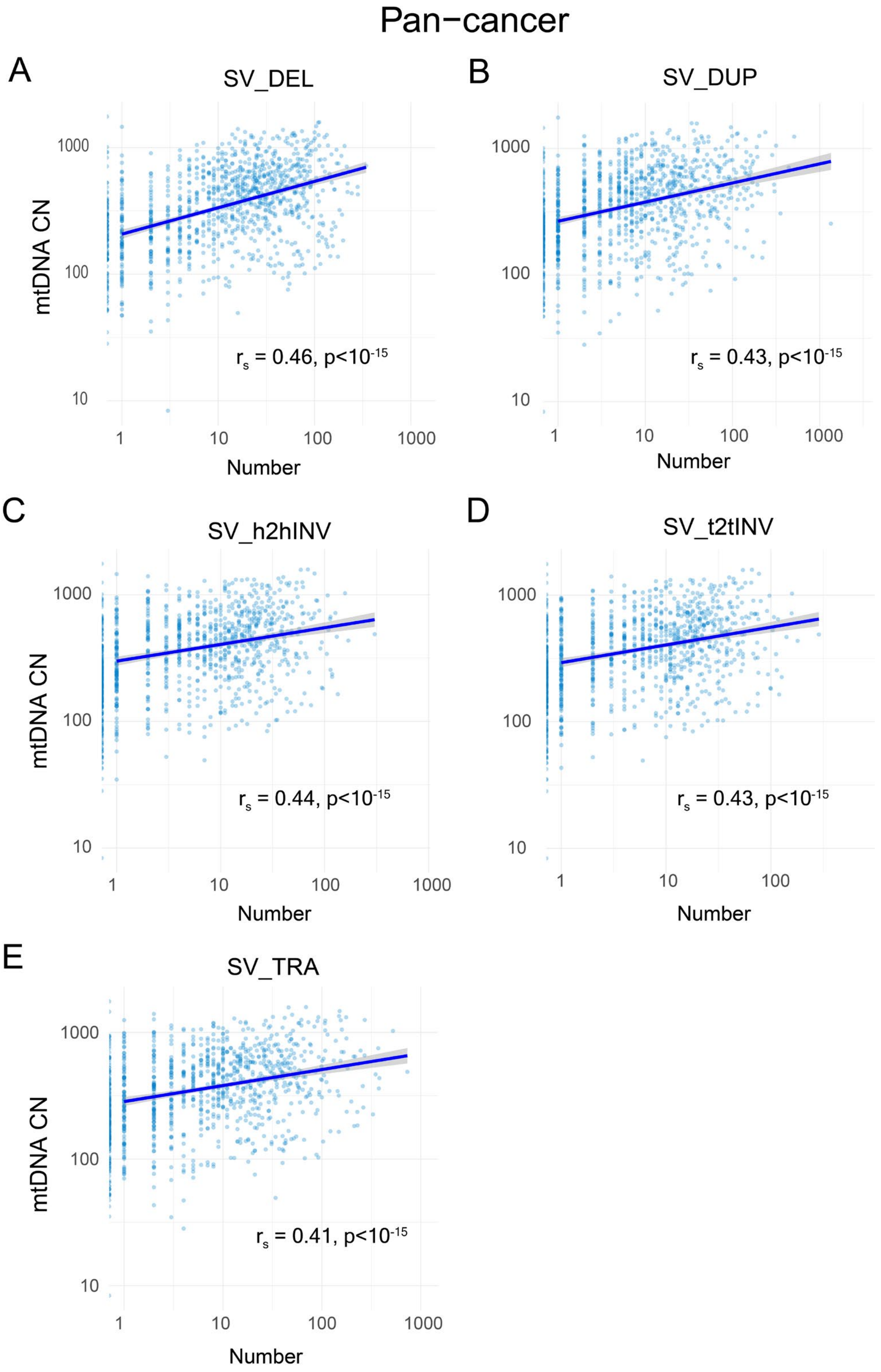


**Figure S18 –** Pan-cancer correlation between mtDNA copy number and the number of structural variants of different types including deletions **(A)**, duplications **(B)**, head-to-head inversions **(C)**, tail-to-tail inversions **(D)** and translocations **(E)**. The solid lines show linear fits to the data and the shaded grey regions show 95% confidence intervals.

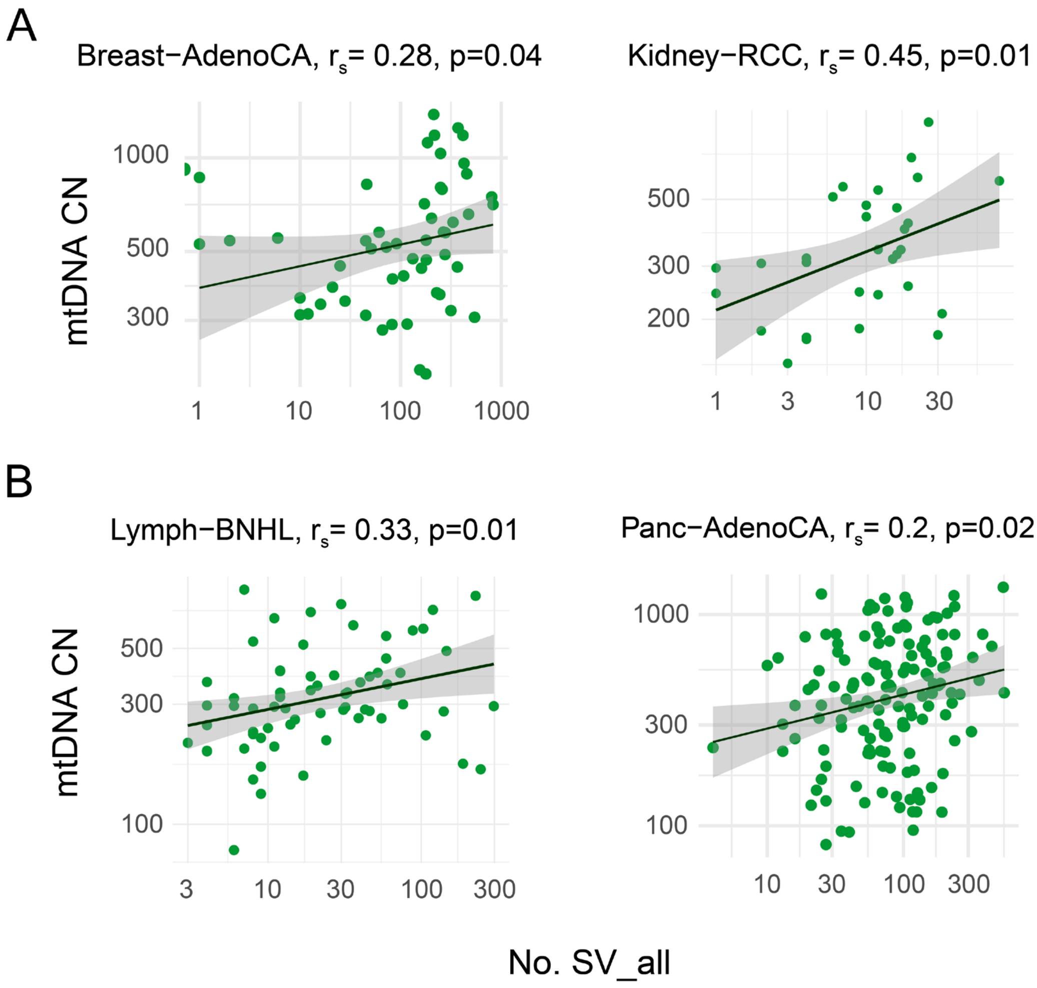


**Figure S19 –** Correlation between mtDNA copy number and the number of structural variants (all types) in individual cancer types. The solid lines show linear fits to the data and the shaded grey regions show 95% confidence intervals.

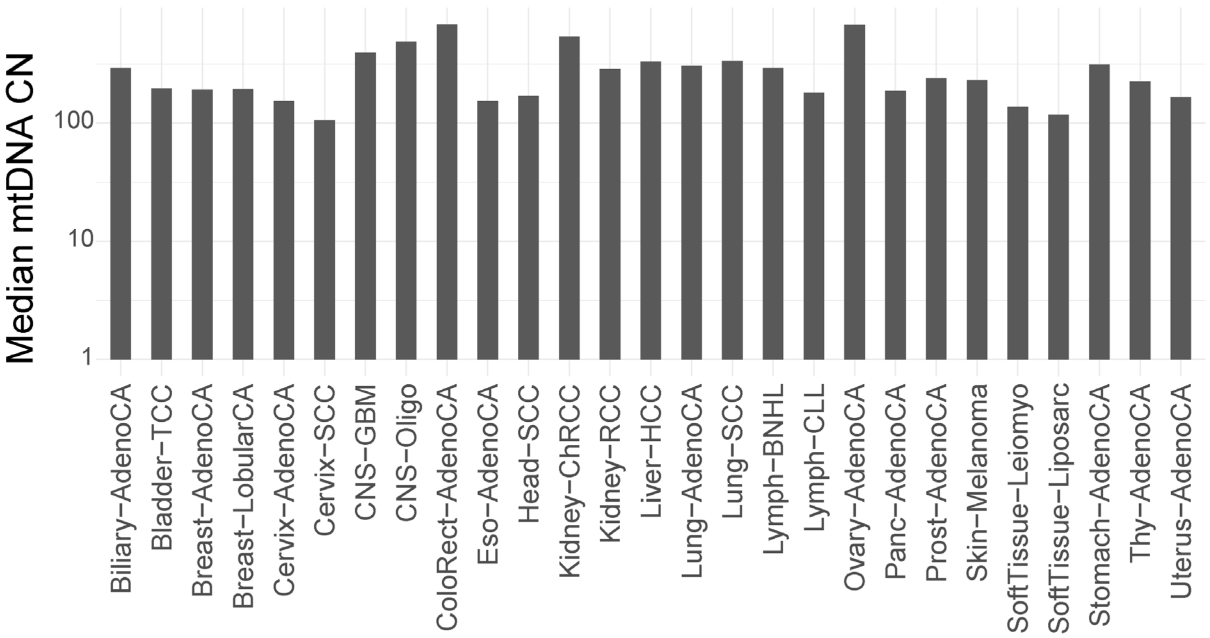


**Figure S20 –** The median mtDNA copy number for each cancer type calculated from the samples included in gene expression analysis.

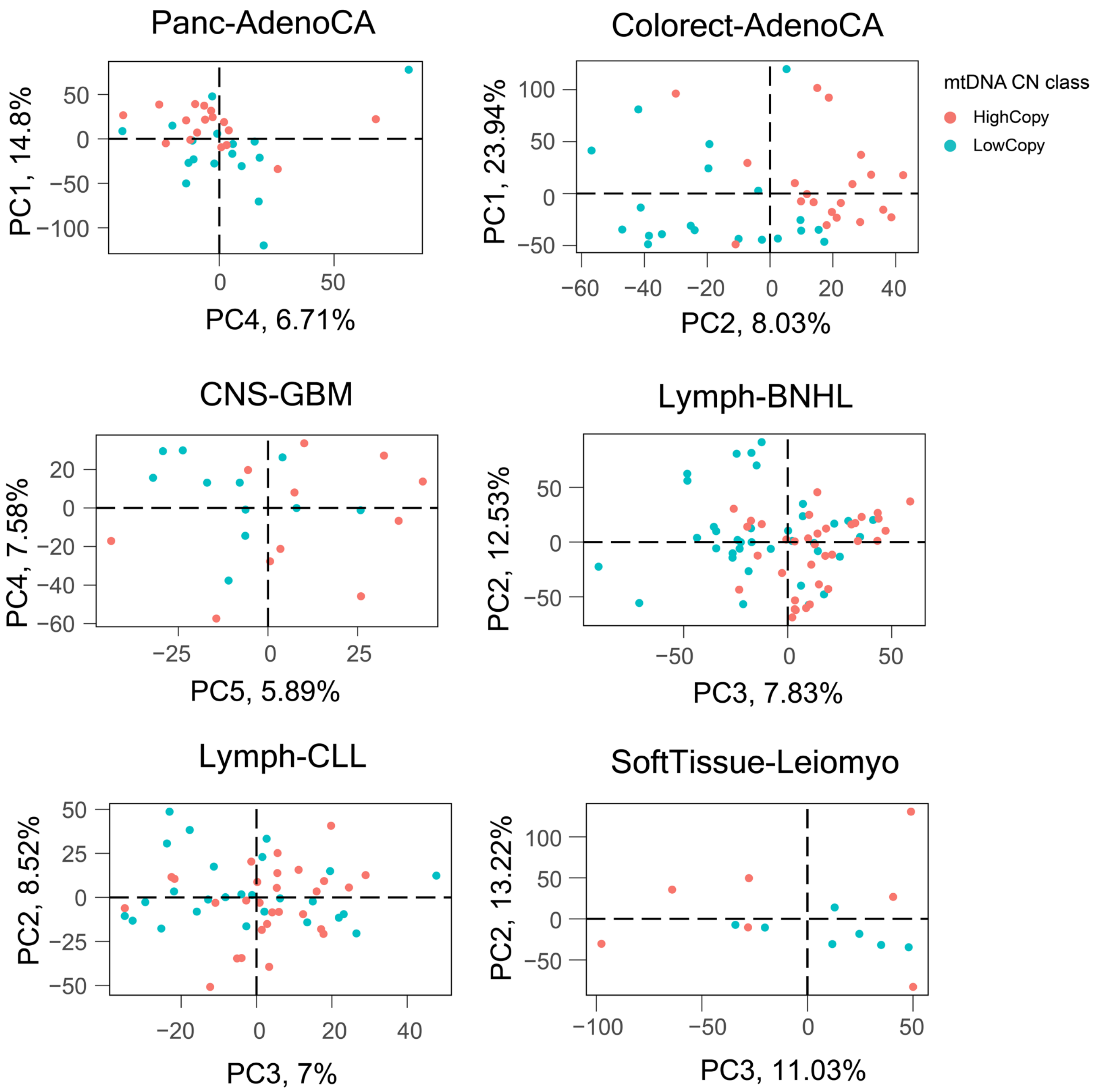


**Figure S21 –** Principal component analysis based on gene expression patterns of high- and low- mtDNA samples in individual cancer types.

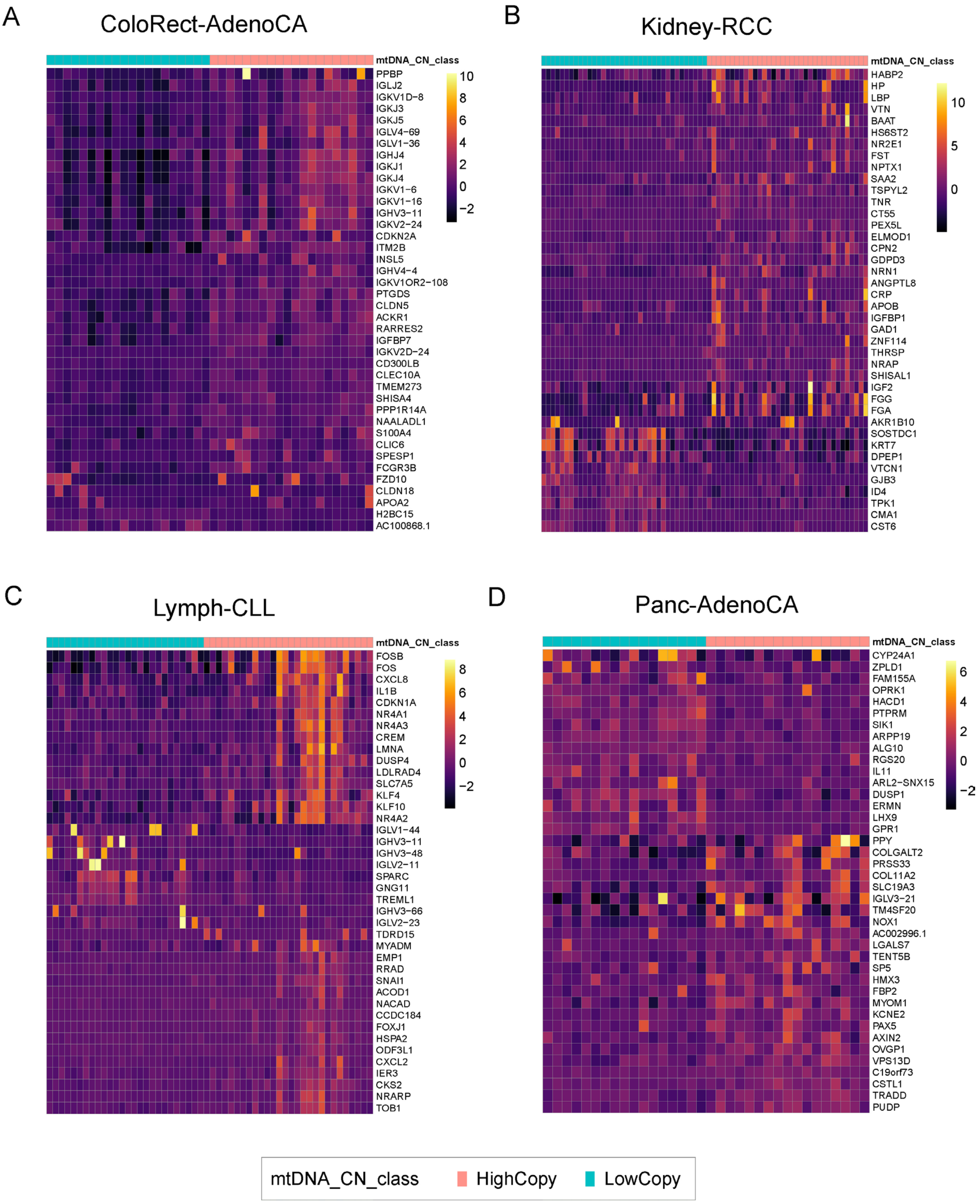


**Figure S22 –** Expression pattern of top differentially expressed genes between high- and low- mtDNA samples in ColoRect-AdenoCA **(A)**, Kidney-RCC **(B)**, Lymph-CLL **(C)**, and Panc-AdenoCA **(D)**.

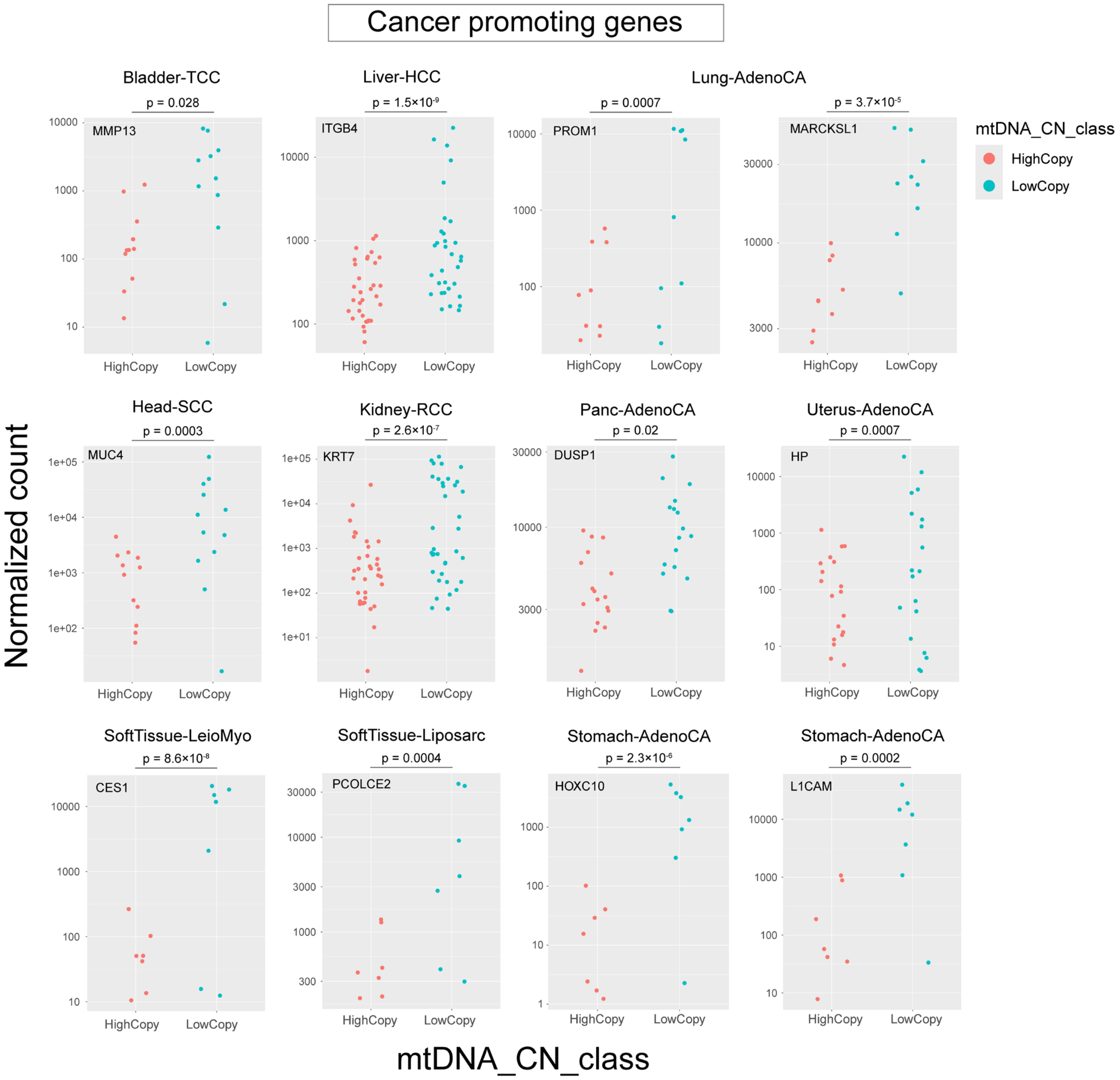


**Figure S23 –** Normalized count values of some cancer promoting genes in high- and low- mtDNA samples in individual cancer types. The p-values show p-adjusted values obtained after FDR correction in DESeq2.

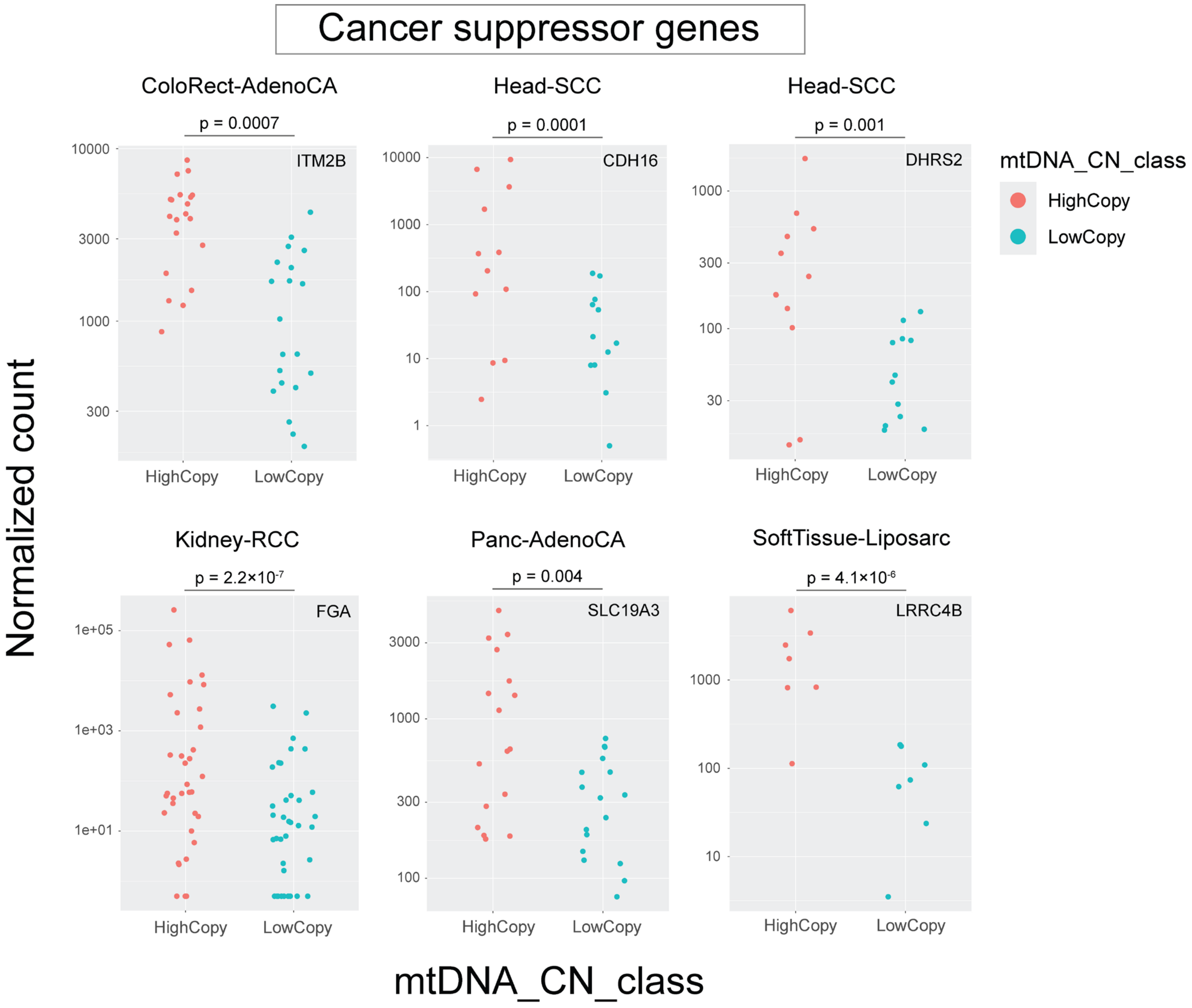


**Figure S24 –** Normalized count values of some cancer suppressor genes in high- and low- mtDNA samples in individual cancer types. The p-values show p-adjusted values obtained after FDR correction in DESeq2.

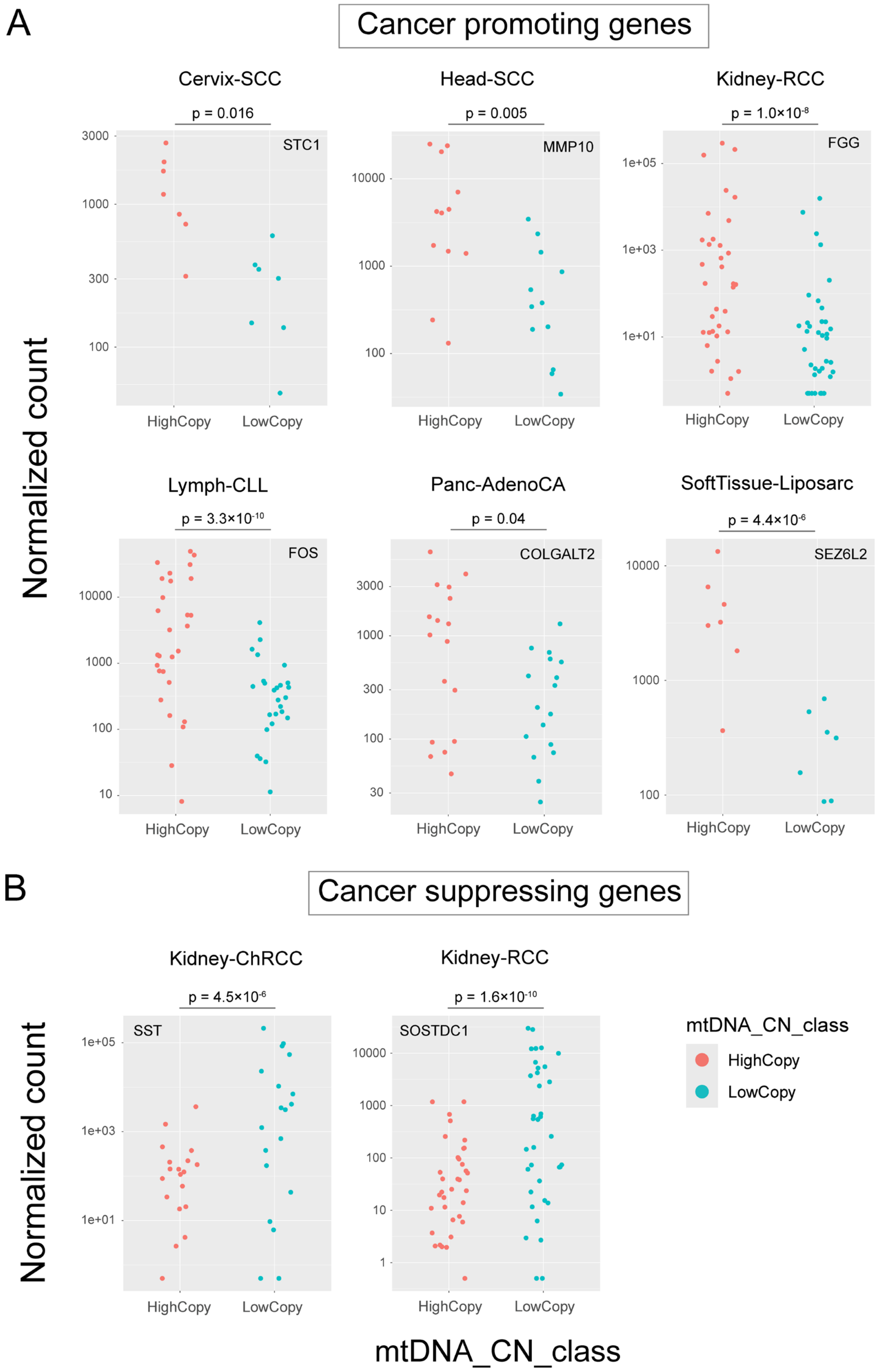


**Figure S25 –** Normalized count values of some cancer promoting and cancer suppressor genes in high- and low- mtDNA samples in individual cancer types. The p-values show p-adjusted values obtained after FDR correction in DESeq2. These cancer promoting genes showed higher expression in high mtDNA samples and these cancer suppressor genes showed higher expression in low mtDNA samples.

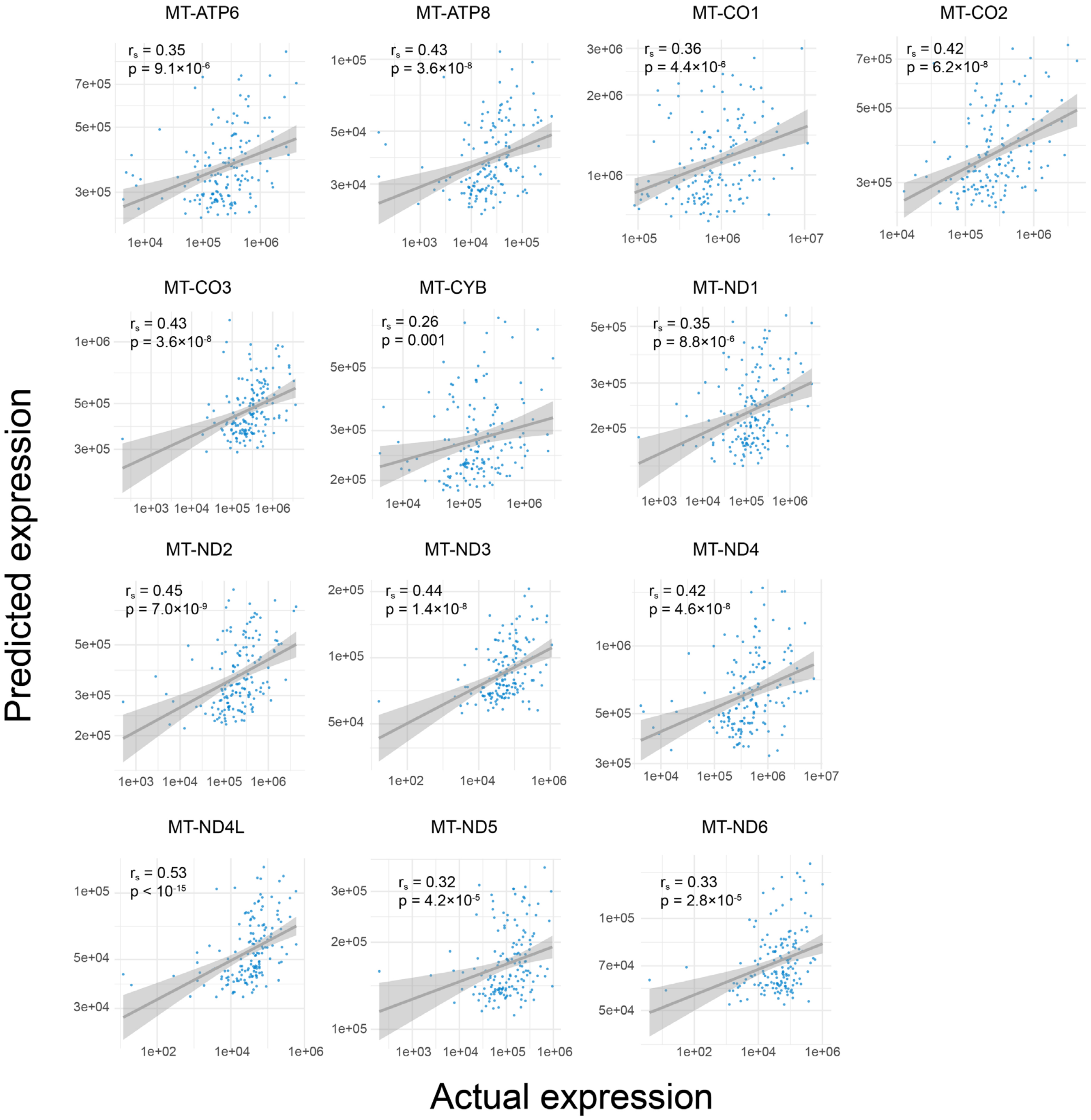


**Figure S26 –** Correlation between actual expression of mtDNA encoded genes and their predicted expression levels from linear regression models with mtDNA copy number as the predictor variable. The solid lines show linear fits to the data and the shaded grey regions show 95% confidence intervals.

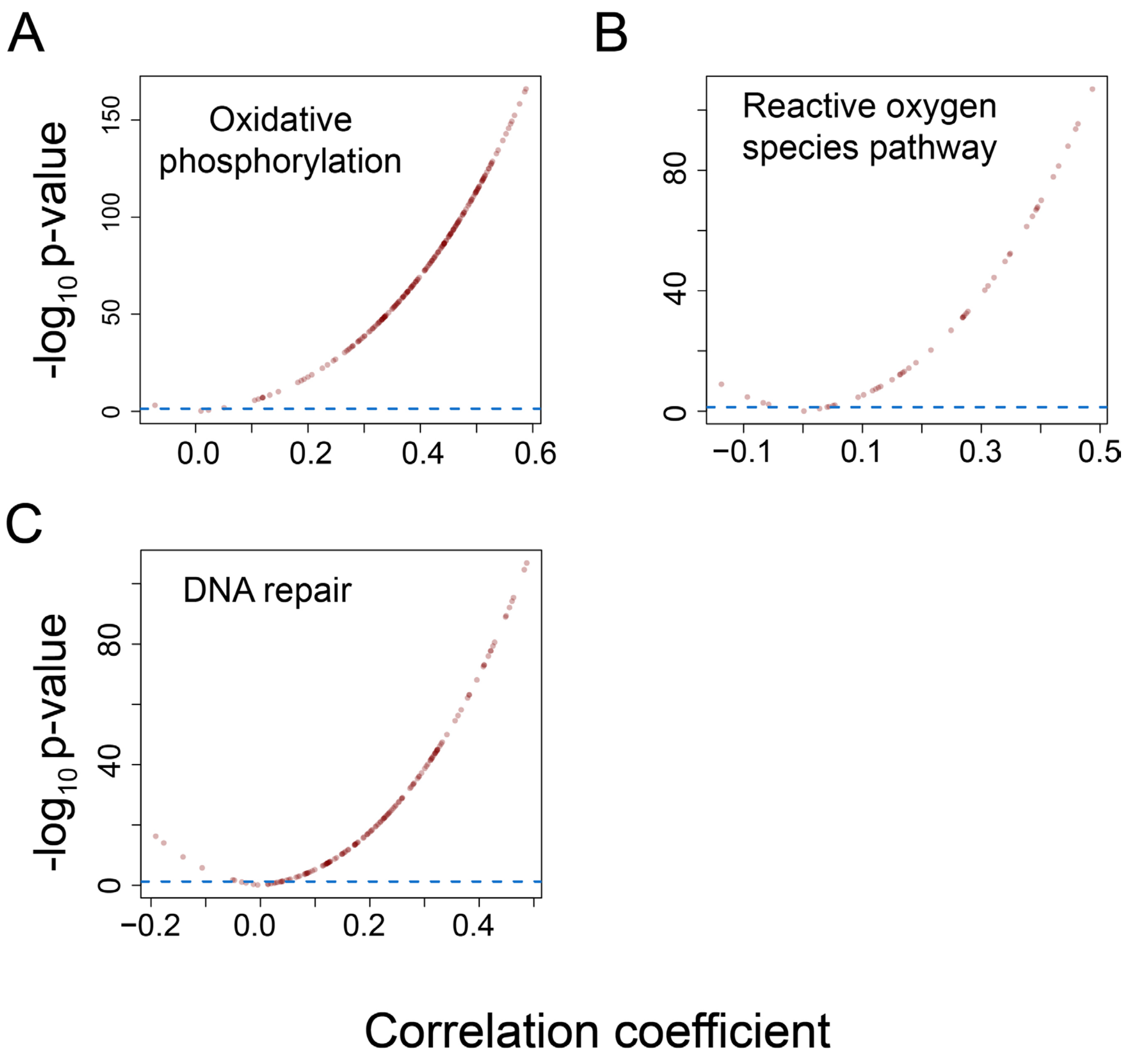


**Figure S27 –** Distribution of Spearman's correlation coefficient and -$\log_{10}$(p-value) of correlations calculated between mtDNA encoded *MT-CO2* gene and the genes in the hallmark gene set in MSigDB associated with oxidative phosphorylation **(A)**, reactive oxygen species **(B)** and DNA repair **(C)**.

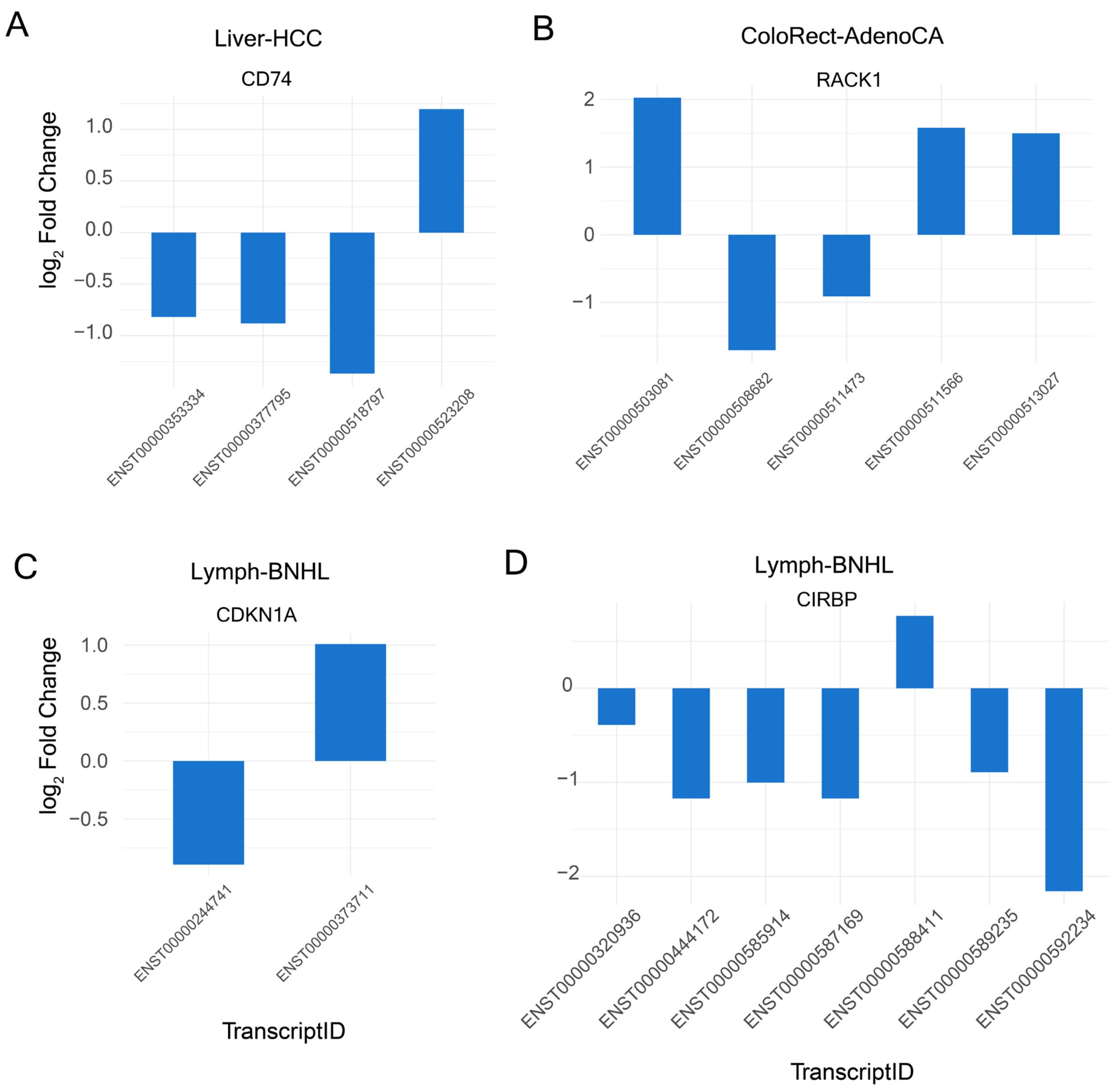


**Figure S28 –** Examples of alternative splicing in specific genes in individual cancer types.

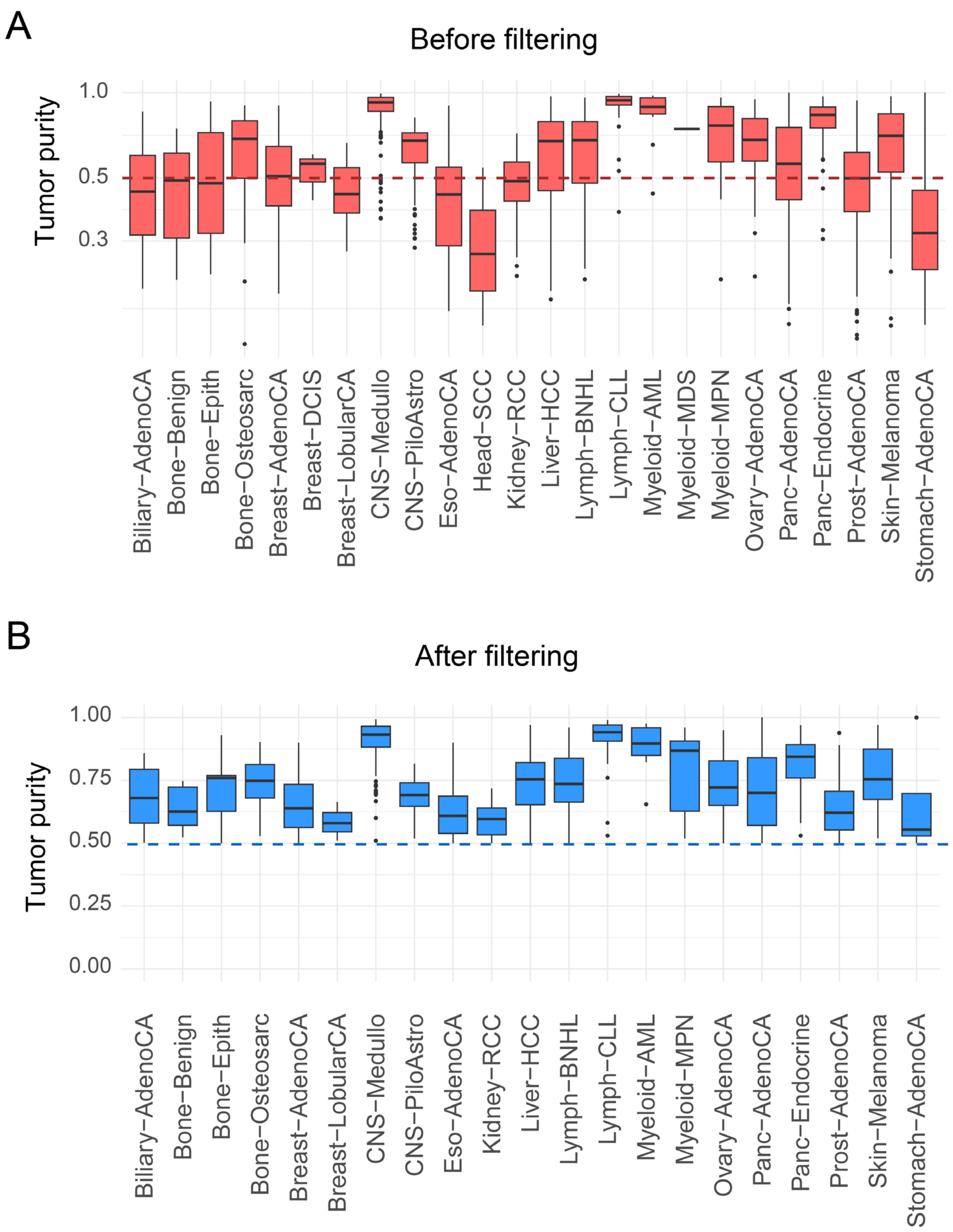


**Figure S29 –** Distribution of tumor purity in individual cancer types before purity filtering **(A)** and after purity filtering **(B)**.

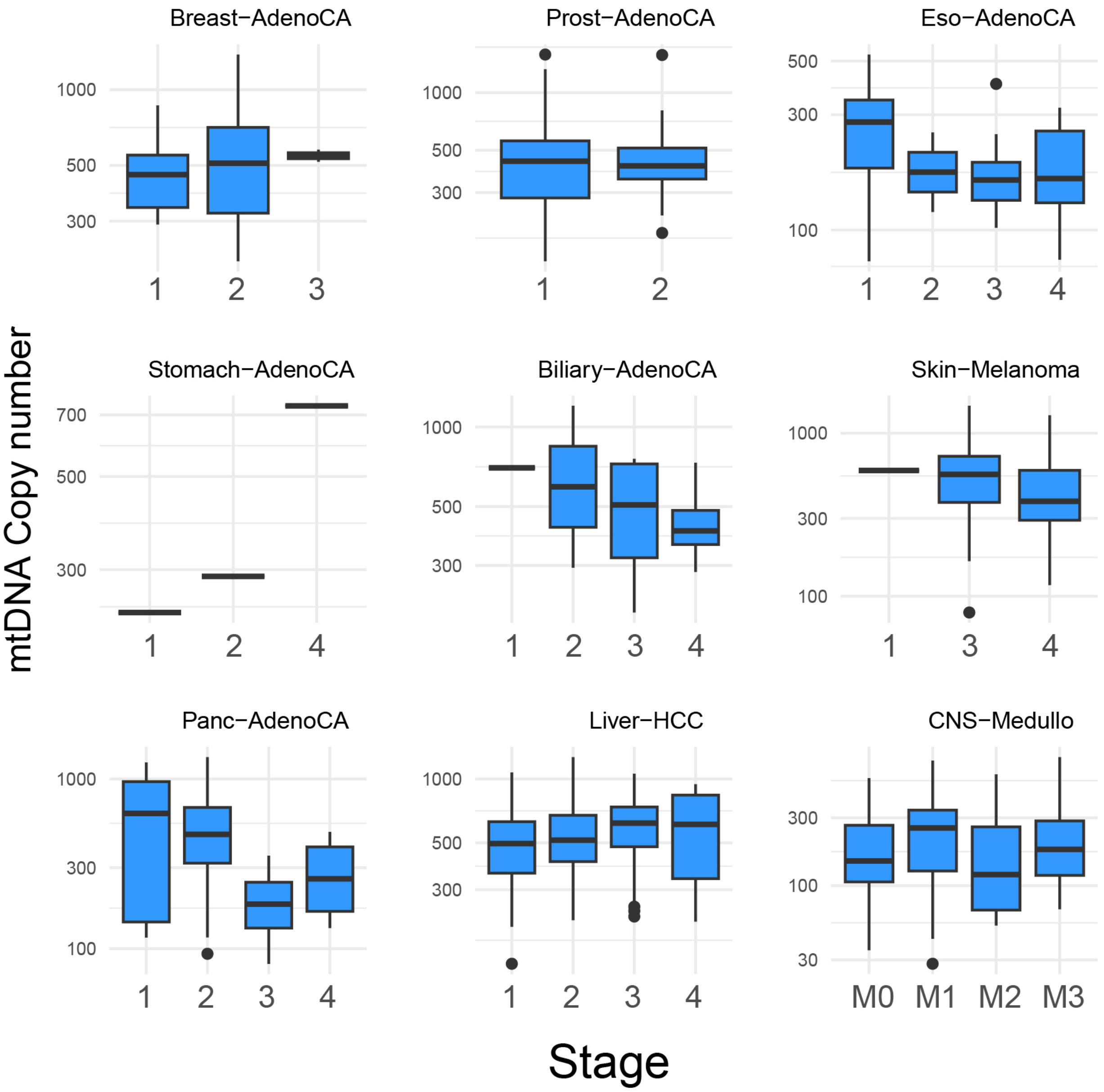


**Figure S30 –** Distribution of mtDNA copy number in samples of different stages in individual cancer types.

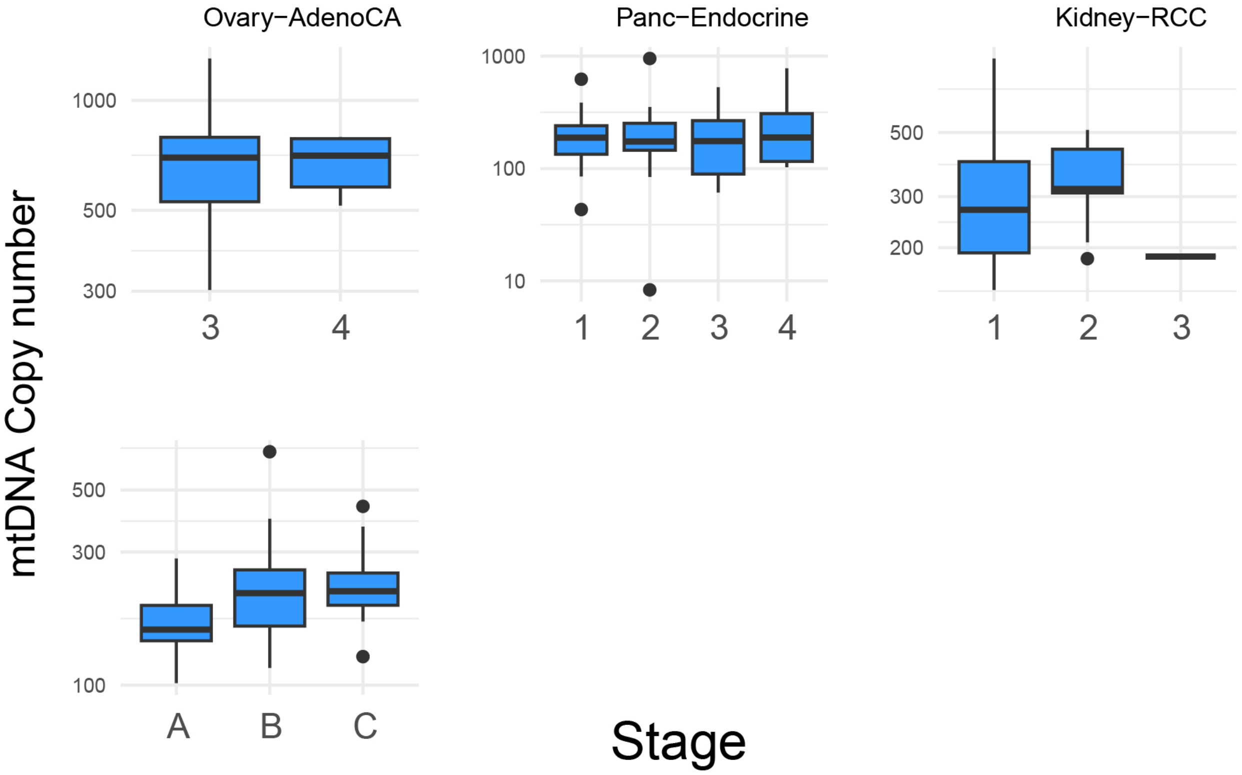


**Figure S30 (continued)**

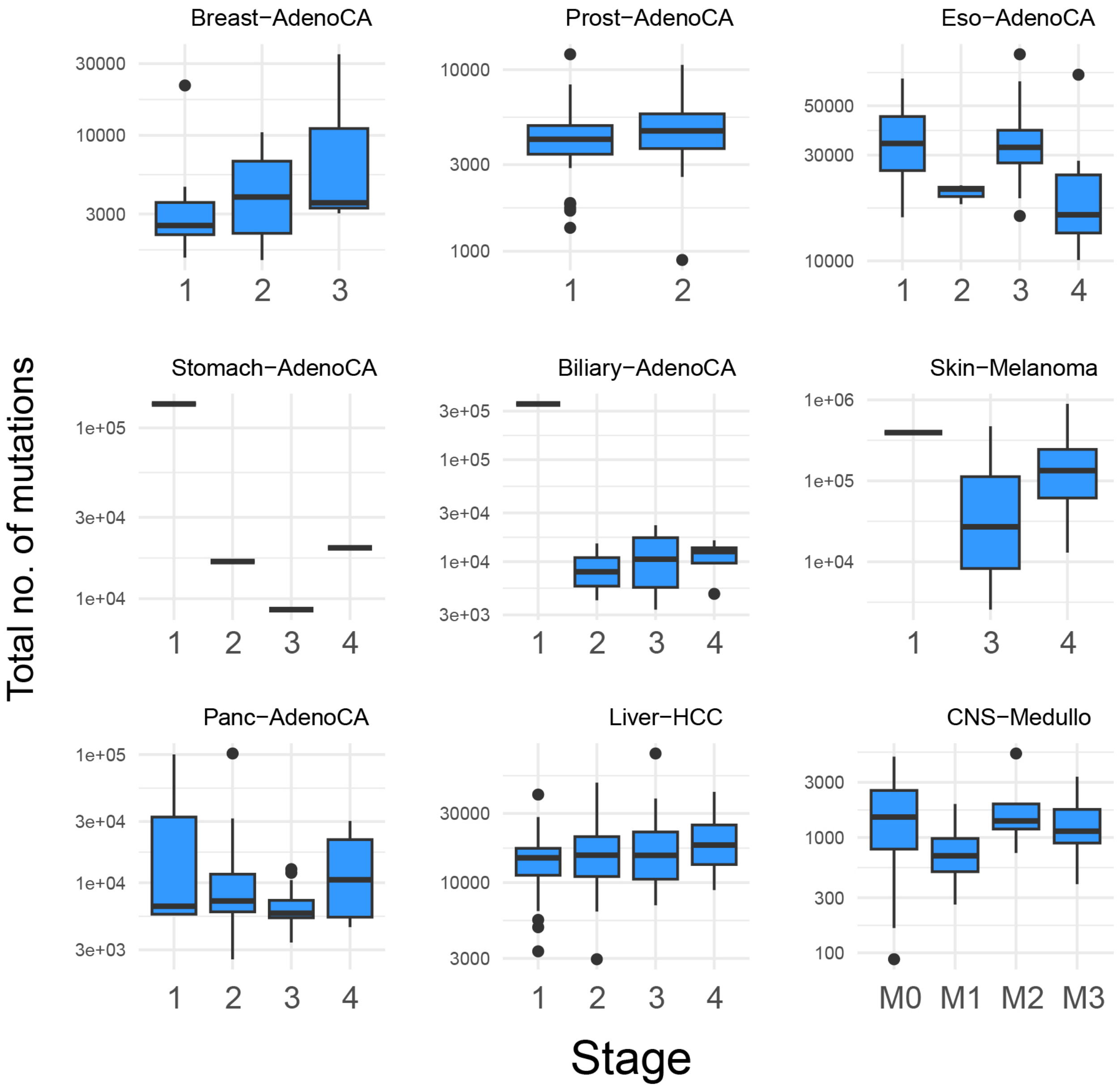


**Figure S31 –** Distribution of total number of mutations in samples of different stages in individual cancer types.

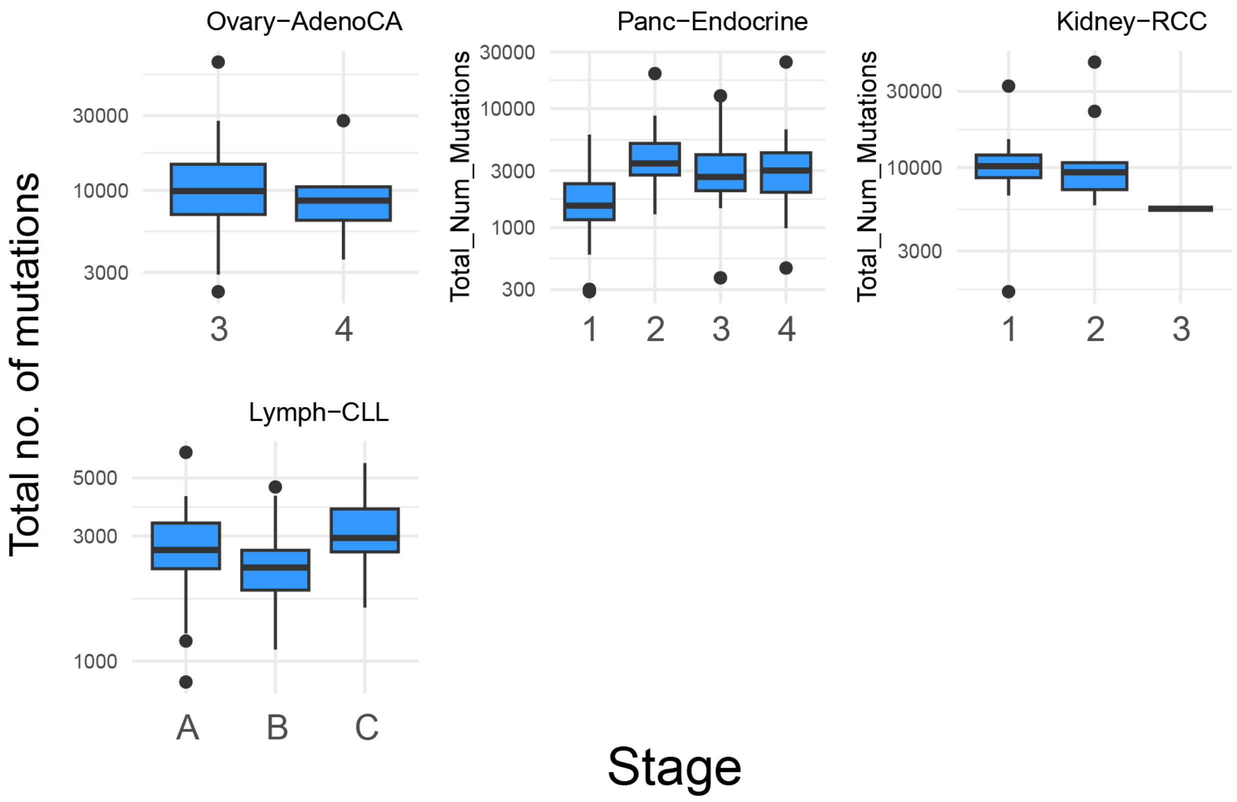


**Figure S31 (continued)**
